# Single-cell multi-omics reveals tumor microenvironment factors underlying poor immunotherapy responses in ALK-positive lung cancer

**DOI:** 10.1101/2024.09.24.614708

**Authors:** Seungbyn Baek, Euijeong Sung, Gamin Kim, Min Hee Hong, Chang Young Lee, Hyo Sup Shim, Seong Yong Park, Hye Ryun Kim, Insuk Lee

## Abstract

Anaplastic lymphoma kinase (ALK) rearrangement is a major oncogenic driver in non-small cell lung cancer (NSCLC). While ALK tyrosine kinase inhibitors have shown promising therapeutic effects, overcoming resistance with immunotherapy becomes necessary when resistance develops. However, various clinical trials have revealed that their efficacies remain limited. To investigate the tumor microenvironment (TME) factors contributing to poor immune checkpoint blockade responses in ALK-positive patients, we performed single-cell RNA and ATAC sequencing on lung adenocarcinoma (LUAD) tumors with and without ALK rearrangements. Integrative analysis with additional public LUAD cohorts revealed distinct immune landscapes in ALK-positive tumors, marked by enriched innate immunity and depleted adaptive immunity. ALK-positive malignant cells exhibit higher stemness and aggressive phenotype. Tumor-associated macrophages (TAMs) in these tumors predominantly maintain pro-tumoral M2-like states, reinforcing immune suppression. B cells show reduced immune reactivity and impaired tertiary lymphoid structure formation, while CD8^+^ T cells display bystander-like signatures and reduced tumor reactivity. Single-cell chromatin accessibility profiles combined with regulatory network analysis suggest that differences in transcription factor activities, rather than chromatin accessibility, may underlie T cell dysfunction. These findings provide insights into the immunosuppressive TME of ALK-positive LUAD, potentially explaining the failure of recent immunotherapy trials and highlighting targets for improving efficacy.

## Introduction

Lung cancer is the leading cause of death by cancer as of 2024, with approximately 80% of cases identified as non-small cell lung cancer (NSCLC)^1^. The identification of multiple driver oncogenic aberrations such as epidermal growth factor receptor (EGFR), Kirsten rat sarcoma (KRAS), anaplastic lymphoma kinase (ALK) and v-raf murine sarcoma viral oncogene homolog B1 (BRAF) in NSCLC has led to the introduction of targeted agents^2^. These therapies have significantly advanced the survival outcomes for lung cancer patients, marking a remarkable improvement in treatment efficacy.

ALK rearrangement occurs in around 5% of NSCLC cases and is more commonly seen in younger, non-smoking patients^3^. ALK-positive lung cancer patients are typically treated with ALK tyrosine kinase inhibitors (TKIs), with fourth-generation TKIs currently in development^4^. However, these inhibitors often lead to the development of resistance, both on-target and off-target. Consequently, there have been numerous efforts to use immunotherapy, specifically anti-PD-1 and PD-L1 blockade, for patients with advanced or resistant ALK-positive NSCLC. Despite these efforts and the observed up-regulation of PD-L1 in ALK-positive tumor cells, clinical trials involving immune-checkpoint blockade (ICB) treatments have not yielded clinical benefits^5^.

Limited studies suggested that the ineffectiveness of ICB treatments might be attributed to factors such as an immunosuppressive tumor microenvironment (TME), impaired T cell functions, and the role of pro-tumoral macrophages^5–7^. A comprehensive study to characterize the TMEs of ALK-positive NSCLCs is needed to better understand the limitations of immunotherapy and to develop potential strategies for improving treatment efficacy. In this study, we generated and collected both single-cell RNA sequencing (scRNA-seq) and single-cell Assay for Transposase Accessible Chromatin with high-throughput sequencing (scATAC-seq) datasets from lung adenocarcinoma (LUAD) patients with ALK rearrangement and those without major oncogenic drivers including EGFR mutation, ALK and ROS1 rearrangement. By thoroughly comparing the TMEs of each mutational status, we aimed to identify characteristics of cancer and immune cells that might elucidate the inadequate responses to immunotherapy.

## Results

### Single-cell multi-omics reveals ALK-driven changes in tumor immune landscapes

To identify characteristics of TMEs that could affect responses to immunotherapy for patients with ALK rearrangement, we generated scRNA-seq and scATAC-seq datasets from patients with ALK rearrangement (ALK) and patients without any major driver mutations including EGFR, ALK, and ROS1 mutations (WT) prior to clinical treatment. Furthermore, we collected additional scRNA-seq datasets from three public cohorts that included ALK-positive LUAD patients^8–10^ **(Supplementary Table 1-2)**. With these single-cell multi-omics datasets, we conducted data integration and several downstream analyses comparing ALK and WT TMEs **(Fig. 1A)**.

**Fig. 1.**
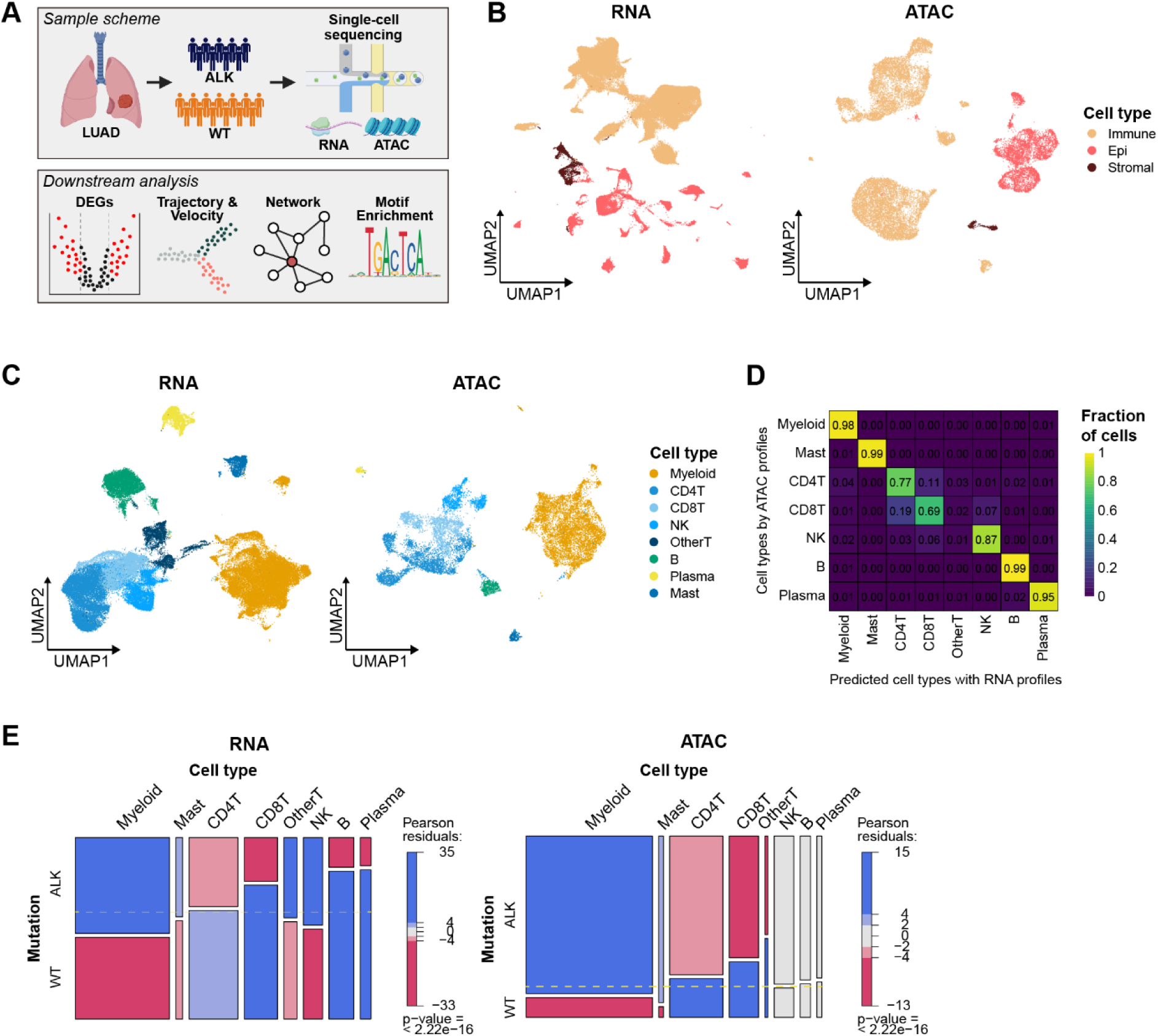
Single-cell multi-omics study for lung adenocarcinoma patients with ALK rearrangement. **A,** Study scheme for single cell multi-omics data and downstream analysis. **B,** UMAP plots of all cells with merged single-cell RNA sequencing (scRNA-seq) profiles (left) and merged single-cell ATAC sequencing (scATAC-seq) profiles (right), color-codes for cell types. **C,** UMAP plots of immune cells with integrated scRNA-seq profiles (left) and integrated scATAC-seq profiles (right), color-codes for cell types. **D,** Heatmap depicting fractions of cell types predicted by scRNA-seq and scATAC-seq profiles. **E,** Mosaic plots for comparison of cell compositions between WT and ALK-positive groups for each omics approach. The color indicates Pearson residuals, showing enrichment or depletion of the cell types for each group. Significance was calculated using the chi-square test.

First, we pre-processed each dataset with ambient RNA removal, scRNA-seq and scATAC-seq data quality controls, and doublet removal. We then merged all datasets from each omics to first identify major TME compartments such as immune, epithelial, and stromal cells **(Fig. 1B)**. We further subclustered cells from the immune compartment and integrated them through batch correction methods to initially inspect palpable differences in immune mechanisms. We identified major immune cell types such as myeloid cells, T cells, NK cells, and B cells for both scRNA-seq and scATAC-seq datasets **(Fig. 1C, Supplementary Table 3)**. For scRNA-seq datasets, while there is dominance in cell counts for in-house cohort (Sev), we observed inclusion of cells with ALK and WT status for each major immune cell type across at least three different cohorts **(Supplementary Fig. 1A)**. Two datasets from Maynard et al. had low overall cell counts which were mostly epithelial cells, which resulted in exclusion of Maynard cohort for some immune cell types. For scATAC-seq datasets, we observed even distribution of cells from both mutation status **(Supplementary Fig. 1B)**. To check concordance of cell type annotations among both single-cell omics datasets, we conducted multi-modal integration and cell type label transfers with scRNA-seq and scATAC-seq datasets. We confirmed that the cells from each major immune cell types were clustered together **(Supplementary Fig. 1C)**. Cell type annotation of scATAC-seq cells with label transfer method using scRNA-seq cells as reference also showed high coherence between cell types from manual annotation and cross-omics reference-based annotation **(Fig. 1D, Supplementary Fig. 1D)**.

To determine which immune cell types were more prevalent in each TME based on mutation status, we performed cell compositional analysis with the immune cells. In ALK-positive samples, there was an enrichment of immune cells associated with innate immunity, such as myeloid cells and NK cells. In contrast, cells involved in adaptive immunity, like T cells and B cells, were found to be depleted **(Fig. 1E)**. These changes in immune compartment by ALK rearrangement suggest insufficient anti-tumor responses due to impaired T cell activation^7^. This conclusion was further corroborated by similar patterns observed in the compositional analysis conducted using scATAC-seq datasets and integrated multi-omics datasets **(Fig. 1E, Supplementary Fig. 1E)**.

### ALK-positive malignant cells exhibit higher stemness and aggressive phenotype

Oncogenic mutations may have more direct and profound impact on epithelial cells. Therefore, we first investigated cellular and molecular changes in epithelial cells by ALK rearrangement. We isolated all epithelial cells and classified them into normal and malignant cells based on inferred copy number variations and reference-based identifications. We observed high patient heterogeneity among malignant cells (**Fig. 2A, Supplementary Fig. 2A**). Furthermore, we observed even greater separations among cells from different patients after isolating only the malignant cells **(Fig. 2B, Supplementary Fig. 2B)**. For preliminary verification of ALK-positive tumor cells, we measured proportions of *ALK* and *PD-L1* expressing malignant cells. As expected, almost all *ALK* expressing malignant cells were from ALK-positive tumors and *PD-L1* expression was upregulated as well^5^ (**Fig. 2C**).

**Fig. 2.**
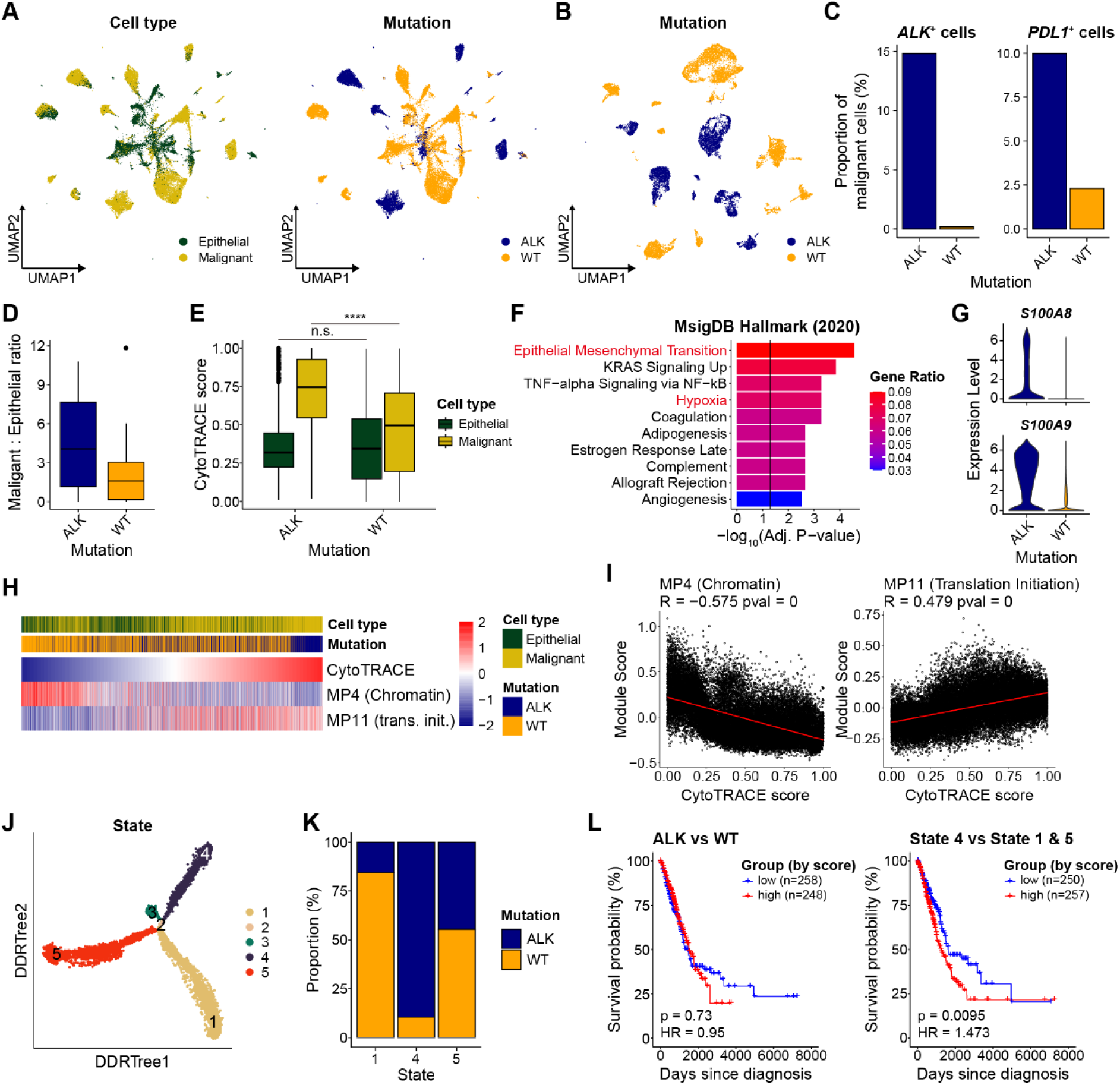
Characterization of ALK-positive malignant cells. **A,** UMAP plots of normal and malignant epithelial cells, color-coded for epithelial cell type (left) or ALK mutation status (right). **B,** UMAP plots of malignant epithelial cells with color codes for ALK mutation status. **C,** Fractions of cells with *ALK* expression (left) or *PDL1* expression (right) within malignant cells for each mutation group. **D,** Ratio of malignant to normal epithelial cells for each mutation group. **E,** Difference of CytoTRACE (tumor plasticity) scores between epithelial cell types for each mutation group. Significance was calculated using two-sided Wilcoxon rank-sum test. n.s. = not significant, **** < 0.0001. **F,** Enrichment of top 100 up-regulated genes in ALK-positive malignant cells compared to WT malignant cells for MsigDB Hallmark (2020) gene sets. The color indicates fractions of the query genes that overlap with each reference gene set. Significance was calculated using Fisher’s exact test with the Benjamini-Hochberg correction. **G,** Violin plots for expression of *S100A8* and *S100A9* in malignant cells for each mutation group. **H,** Heatmaps for cells sorted by the scaled CytoTRACE scores. Each cell’s gene signature scores for each meta-program (MP4 and MP11), cell type, and mutation group are indicated. **I,** Dot plots of CytoTRACE scores and module scores for MP4 and MP11 with a line for their linear regression. P-values and coefficients for correlation were calculated with two-sided Pearson correlation test. **J,** Trajectory plot with malignant cells with DDRTree algorithm. The cells are colored with their states or branches along the trajectory. **K,** Proportions of ALK-positive or WT cells for each state. Proportions are fixed by total malignant cell counts for each mutation. **L,** Keplan-Meier survival plots with TCGA LUAD samples grouped with high or low gene scores for DEGs between ALK vs WT group (left) and DEGs between State 4 vs State 1 and 5 groups (right). Significance was calculated using log-rank test and hazard ratio (HR) was calculated using Cox regression.

Next, we wanted to determine whether ALK-positive malignant cells would have more aggressive phenotypes. Aggressive cancer exhibits a sustained increase of malignant cell proliferation driven by oncogenic signaling^11^. Therefore, we measured ratios between malignant cells and normal epithelial cells for each sample. ALK-positive tumors had four times more malignant cells on average while WT tumors had about 1.5 times more malignant cells **(Fig. 2D, Supplementary Fig. 2C)**. Furthermore, we estimated plasticity of each epithelial cell using CytoTRACE^12^ to determine their stemness (developmental potential). For normal epithelial cells, there were no differences in stemness between WT and ALK-positive tumors. However, in malignant cells, we observed the significant increases in stemness for ALK-positive tumors **(Fig. 2E**). Through differentially expressed gene (DEG) analysis of malignant cells between WT and ALK-positive samples, we selected the top 100 up-regulated genes in ALK-positive tumor (**Supplementary Table 4**) and found that they were associated with MSigDB hallmark gene set^13^ relevant to metastatic aggressive cancers, such as epithelial mesenchymal transition (EMT) and hypoxia pathways^14^ **(Fig. 2F)**. Furthermore, genes related to immune suppression by tumor cells such as *S100A8* and *S100A9,* which likely contribute to immune evasion, were among the top 100 up-regulated genes in ALK-positive malignant cells^15^ **(Fig. 2G)**.

To dissect heterogeneous cancer cells and their functional states, pan-cancer meta-programs (MPs) were recently defined using numerous single-cell datasets^16^. We assessed gene signature scores for malignant cells with these well-defined MPs relevant to lung cancer cells. In ALK-positive malignant cells, we observed MP enrichment patterns that mirrored those seen in MSigDB hallmark gene set enrichment analyses, showing association with EMT, Stress, and Hypoxia pathways **(Supplementary Fig. 2D)**. Moreover, there was an enrichment of MPs related to interferon and major histocompatibility complex (MHC), suggesting increased interactions with the immune compartment in ALK-positive tumors. When we arranged the malignant cells based on stemness scores, we noted a decrease in chromatin-related MP scores and an increase in translation initiation scores, accompanied by an increase in malignant and ALK-positive cells **(Fig. 2H-I)**. These results suggest aggressive tumor phenotype associated with DNA instability and higher cellular activities in ALK-positive malignant cells.

To identify subsets of malignant cells within tumors that have a higher progression potential, we constructed developmental trajectories with ALK-positive and WT malignant cells **(Fig. 2J)**. Among the major states, state 5 included both ALK-positive and WT malignant cells while state 4 and state 1 were predominantly enriched with ALK-positive malignant cells and WT malignant cells, respectively **(Fig. 2K)**. We observed higher stemness scores for malignant cells of the state 4, the ALK-associated state (**Supplementary Fig. 2E**). State 4 malignant cells also showed higher expression of *S100A8* and *S100A9*, as well as MPs for stress, hypoxia, and EMT, indicating immune suppression by tumor cells (**Supplementary Fig. 2F-G**). These suggest that the patients with tumors enriched with state 4 malignant cells would have worse clinical outcomes regardless of ALK mutation status. To evaluate the prognostic effect of the gene expression signature of state 4, we performed survival analysis on LUAD patients in The Cancer Genome Atlas (TCGA)^17^ using DEGs (q-value < 0.01, top 100 genes from each comparison group) derived from comparing ALK-positive to WT malignant cells within the trajectory and those from comparing state 4 to state 1 and 5 (**Supplementary Table 5**). While DEGs comparing all ALK-positive and WT malignant cells did not distinguish survival probabilities, we found that patients with higher expression of state 4 DEGs and lower expression of state 1 and 5 DEGs had significantly worse survival outcomes (**Fig. 2L**). This suggests that the gene signatures of the ALK-positive malignant cells with a state 4 transcriptomic phenotype may be linked to poor clinical outcomes in general LUAD contexts. In conclusion, ALK-rearrangement for malignant cells would result in more aggressive cancer states with potential for metastasis and immune suppression.

### Macrophages are more pro-tumoral and immune-suppressive in ALK-positive tumors

To identified immune cell types that are most affected by ALK-positive malignant cells, we conducted cell-cell interaction (CCI) analysis between malignant cells and immune cells. We then compared differences in CCIs between ALK-positive and WT malignant cells. While interactions from ALK-positive malignant cells to all immune cell types increased compared to those from WT malignant cells, interactions with myeloid cells showed the greatest increase (**Fig. 3A**). We found that the interactions from malignant cells to myeloid cells increased mainly through the extracellular matrix components such as laminin and collagen of malignant cells and CD44 of myeloid cells **(Fig. 3B)**. Expression of *CD44* for myeloid cells was recently identified to promote tumor progression through M2-like macrophage polarization in the pan-cancer research^18^. Furthermore, MHC class 2 molecules upregulated for ALK-positive malignant cells also increased interactions with myeloid cells through the receptor encoded by *CD4* gene. Therefore, we next examined biological processes affected by mutation status in myeloid compartment.

**Fig. 3.**
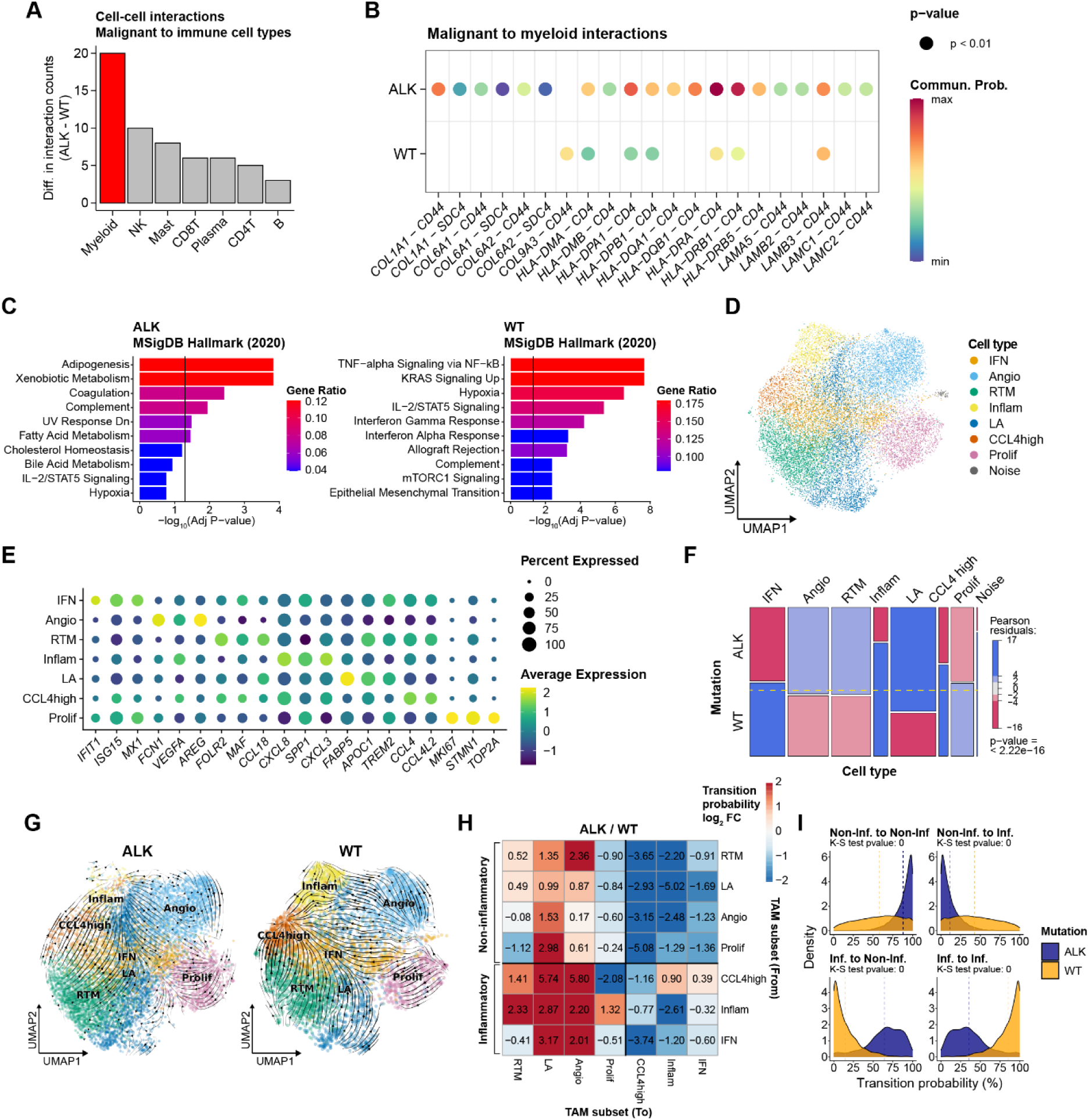
Tumor-associated macrophages (TAMs) in ALK-positive tumors. **A,** Differences in the number of cell-cell interactions from each immune cell type to malignant cells between ALK-positive and WT samples. **B,** Interacting ligand-receptor gene pairs between malignant cells and myeloid cells for each mutation group. Only the significant interactions for at least one group were visualized. **C,** Enrichment of top 50 up-regulated genes in TAMs of ALK-positive tumors (left) and those of WT samples (right) for MsigDB Hallmark (2020) gene sets. The color indicates fractions of query genes that overlap with genes of each reference gene set. The p-values was calculated using the Fisher’s exact test with the Benjamini-Hochberg correction. **D,** UMAP plot for sub cell types of TAMs. **E,** Dot plot for marker genes for each sub cell type of TAMs. **F,** Mosaic plots for comparison of cell compositions between WT and ALK-positive samples for sub cell types of TAMs. The color indicates Pearson residuals, showing enrichment or depletion of the cell types for each mutation group. Significance was calculated using chi-square test. **G,** Stream plots depicting cell transitions according to their RNA velocity for TAMs in ALK-positive tumors (left) and WT tumors (right). **H,** Heatmap depicting cell type transitions from row to column. The color indicates log2 fold changes calculated by dividing transition probability for TAMs in ALK-positive tumors by TAMs in WT tumors. **I,** Transition probabilities among inflammatory TAMs and non-inflammatory TAMs for each mutation group. Significance of differences in distributions of transition probabilities was calculated using Kolmogorov-Smirnov test.

Through sub-clustering of myeloid cells, we identified monocytes, neutrophils, dendritic cells (DCs), and tumor-associated macrophages (TAMs) using their corresponding markers **(Supplementary Fig. 3A-B, Supplementary Table 6)**. Among these myeloid cell subtypes, ALK-positive tumors had significantly higher proportions of TAMs **(Supplementary Fig. 3C)**. This suggests that TAMs could play a significant role in generating pro-tumoral condition of TMEs in ALK-positive tumors.

Using top 50 up-regulated genes for TAMs in ALK-positive tumor and those in WT tumor (**Supplementary Table 7**), we performed enrichment analysis for MSigDB hallmark, Reactome and GO biological process gene sets^13,19,20^. The analysis revealed that TAMs of ALK-positive tumors were associated with M2-like macrophage functions, such as adipogenesis and lipid metabolic processes. In contrast, TAMs of WT tumors were associated with M1-like immune-related functions, including TNF-alpha signaling and interferon gamma response (**Fig. 3C, Supplementary Fig. 3D)**. To further classify functional states of the macrophages, we sub-clustered TAMs and discovered seven TAM sub-states with markers from recent compilation of single-cell-based macrophage markers^21^ **(Fig. 3D-E, Supplementary Fig. 3E, Supplementary Table 8)**. Through compositional analysis, we observed enrichment of inflammation-related TAM states, such as interferon (IFN) TAMs, Inflamed (Inflam.) TAMs, and *CCL4*-high TAMs, in WT tumors, whereas ALK-positive tumors showed enrichment of metabolic TAM states, such as angiogenesis (Angio) TAMs and lipid-associated (LA) TAMs **(Fig. 3F)**.

Because macrophage states are often temporary and there is consistent transition between macrophage states in the TME^22^, we conducted RNA velocity analysis to identify transient or persistent states in TAMs of ALK-positive and WT tumors. We observed the clear trend of transition into inflammation-related TAM states in WT tumors, while ALK-positive tumors showed transition into LA TAMs **(Fig. 3G)**. We then quantified transition probability between TAM states and observed higher persistence for metabolic TAMs in ALK-positive tumors, whereas inflammatory TAMs showed significantly higher persistence in WT tumors **(Supplementary Fig. 3F)**. Direct comparisons between ALK-positive and WT tumors also showed a higher transition probability into metabolic TAMs for ALK-positive tumors and into inflammatory TAMs for WT tumors **(Fig. 3H-I)**. These results collectively suggest that ALK-positive tumors have more M2-like TAMs than WT tumors and that TAMs of ALK-positive tumors exhibit a higher tendency towards metabolic TAM rather than inflammatory TAM states. Consequently, pro-tumoral and immune-suppressive TMEs are maintained through macrophages in ALK-positive tumor.

### B cells in ALK-positive tumors have reduced immune reactivity and tertiary lymphoid structure (TLS) formation capacity

In contrast to innate immunity, adaptive immune compartments were notably depleted in ALK-positive tumors, potentially indicating a reduced capacity to sustain anti-tumoral immune functions^23^. Thus, we first examined B cells for their humoral responses against tumor. Through subclustering analysis, we identified B cell subtypes, including memory B cells, cycling B cells, and plasma cells, based on their marker gene expressions **(Fig. 4A-B, Supplementary Fig. 4A, Supplementary Table 9)**. Both memory B cells and plasma cells were more abundant in WT tumors **(Fig. 4C)**.

**Fig. 4.**
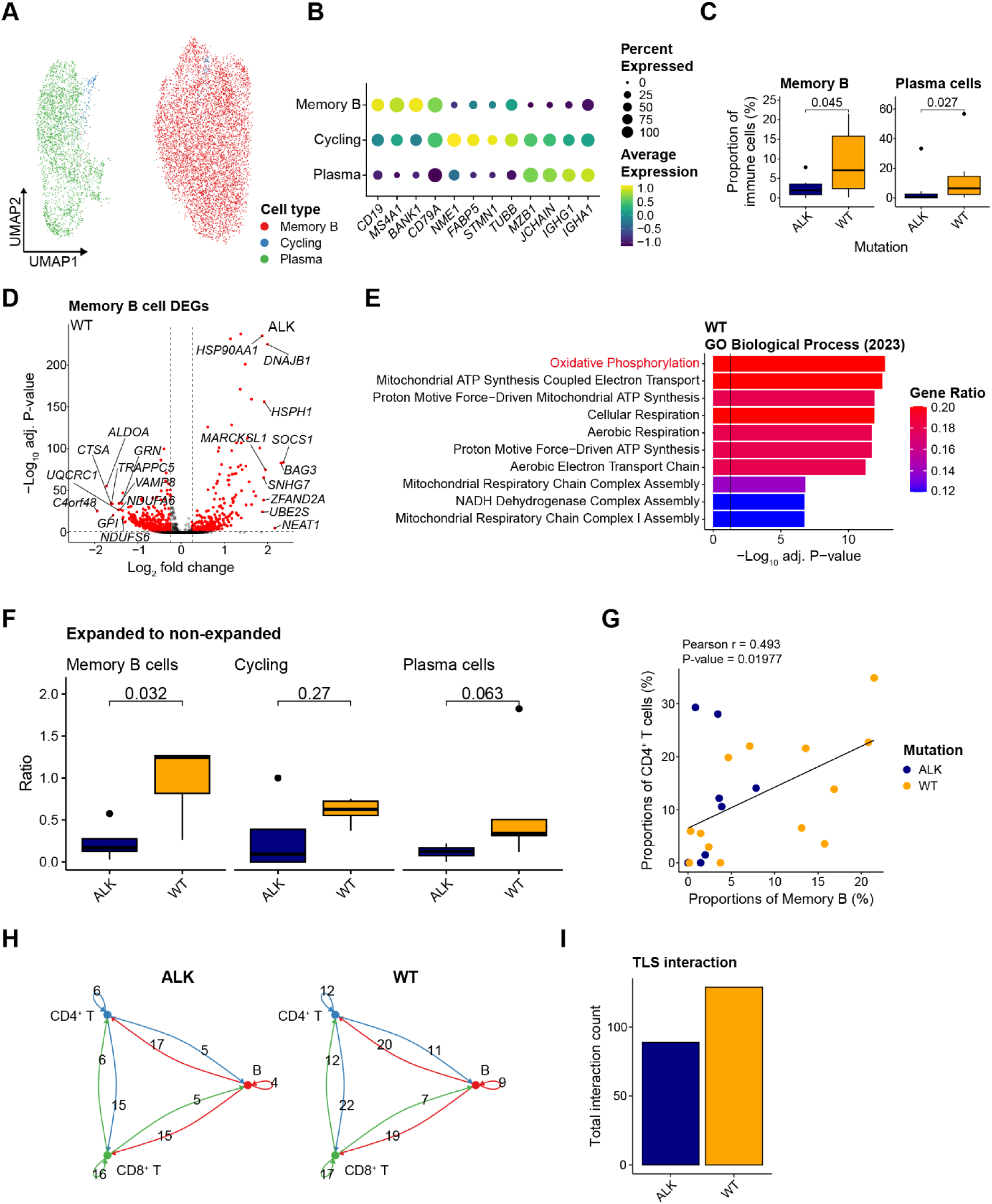
B cells in ALK-positive tumors. **A,** UMAP plots of B cells, color-coded for sub cell types. **B,** Dot plot for marker genes for each sub cell type of B cells. **C,** Proportions of B cells in tumor for each mutation group. Significance of difference was calculated using the two-sided Wilcoxon rank-sum test. **D,** Volcano plot of gene expression differences in B cells between WT and ALK-positive tumors. **E,** Enrichment of top 50 up-regulated genes in memory B cells of WT tumors for GO biological process (2023) gene sets. The color indicates fractions of query genes that overlap with genes from the reference gene sets. The p-values was calculated using the Fisher’s exact test with the Benjamini-Hochberg correction. **F,** Ratio of expanded to non-expanded B cell subtypes for each mutation group. Significance of difference was calculated using the two-sided Wilcoxon rank-sum test. **G**, Scattered plot of relationship between proportion of B cells and proportion of CD4^+^ T cells in each of mutation group. **H**. Cell-cell interaction graph among B cell, CD8^+^ T cells, and CD4^+^ T cells in ALK-positive tumors (left) and WT tumors (right). Edge color corresponds to the sender cell type (red for B cells, green for CD8^+^ T cells, blue for CD4^+^ T cells). **I**. Bar plots depicting total interactions among three cell types for tertiary lymphoid structure (TLS) in ALK-positive and WT tumors.

From DEG analysis of memory B cells between ALK-positive and WT tumors, we identified up-regulation of heat-shock protein genes in ALK-positive tumors (**Fig. 4D**). Additionally, we found that the top 50 up-regulated genes in memory B cells of WT tumors are enriched in oxidative phosphorylation while the top 50 up-regulated genes in memory B cells of ALK-positive tumors are enriched for heat-shock protein-related pathways **(Fig. 4E, Supplementary Fig. 4B)**. The enrichment of oxidative phosphorylation in B cells of WT tumors suggests not only a compositional increase but also heightened functional activation and effector capabilities^24^. Conversely, the expression of heat-shock protein genes in memory B cells of ALK-positive tumors suggests a stress-responsive state, which has recently been linked to unfavorable outcomes for ICB responses^25^.

We reconstructed B cell receptor (BCR) sequences from scRNA-seq data to analyze B cell expansion status. We observed a significantly lower ratio of expanded to non-expanded memory B cells in ALK-positive tumors compared to WT tumors **(Fig. 4F)**. These findings suggest that B cells in ALK-positive tumors have a reduced capability for adaptive anti-tumoral responses.

Since B cells are crucial components of TLSs and the formation of TLSs is vital for immunotherapy responses^26^, we examined potential differences in TLS-like structure components between ALK-positive and WT tumors. Positive correlations were found between the proportions of B cells and CD4^+^ T cells in immune environments, with WT tumors generally having more B cells **(Fig. 4G)**. Reduced interactions among B cells and T cells in WT environments were confirmed, suggesting less formation of TLS-like structures in the ALK-positive tumors **(Fig. 4H-I)**. Overall, differences in compositional enrichment, activation status, and interactions among components of TLS-like structures indicated inadequate anti-tumoral responses by B cells of ALK-positive tumors which could possibly lead to unfavorable TMEs for immunotherapy responses.

### CD8^+^ T cells in ALK-positive tumors exhibit reduced tumor-reactivity and increased bystander traits

Next, we investigated the impact of ALK rearrangement on CD8^+^ T cells due to their direct involvement in ICB treatment and responses through anti-tumoral activities^27^. For favorable ICB responses, the presence of tumor-reactive cytotoxic CD8^+^ T cells that can be reinvigorated is necessary^28^. Through sub-clustering analysis, we identified subtypes of CD8^+^ T cells using specific marker genes: naïve cells expressing *IL7R*, memory cells expressing *GZMK* and *CCR7*, tissue-resident-like IFNG-high cells expressing *CD69* and *IFNG*, stress-responsive heat-shock protein (HSP) high cells, IFN high cells expressing interferon-related genes, effector cells expressing *GZMB* and *PRF1*, and exhausted cells expressing effector signature genes along with exhaustion markers such as *PDCD1* and *CXCL13* **(Fig. 5A-B, Supplementary Fig. 5A, Supplementary Table 10)**. Through compositional analysis, we observed a depletion of exhausted CD8^+^ T cells for ALK-positive tumors **(Fig. 5C)**. Therefore, we conducted DEG analysis to identify underlying molecular changes in CD8^+^ T cells in tumor-reactive but possibly dysfunctional states. Interestingly, genes conventionally related to T cell exhaustion, such as *CXCL13*, *TIGIT*, and *HAVCR2* were enriched in CD8^+^ T cells of WT tumors, while NK receptors like *KLRK1* and *KLRC2,* along with NK cell markers such as *XCL1,* were enriched in CD8^+^ T cells of ALK-positive tumors (**Fig. 5D**). Given that *ENTPD1* and *ITGAE* indicate tumor reactivity^29^, while NK receptors often indicate bystander T cells^30^, these results imply that these ‘exhausted’ CD8^+^ T cells in ALK-positive tumors became dysfunctional through noncanonical mechanisms.

**Fig. 5.**
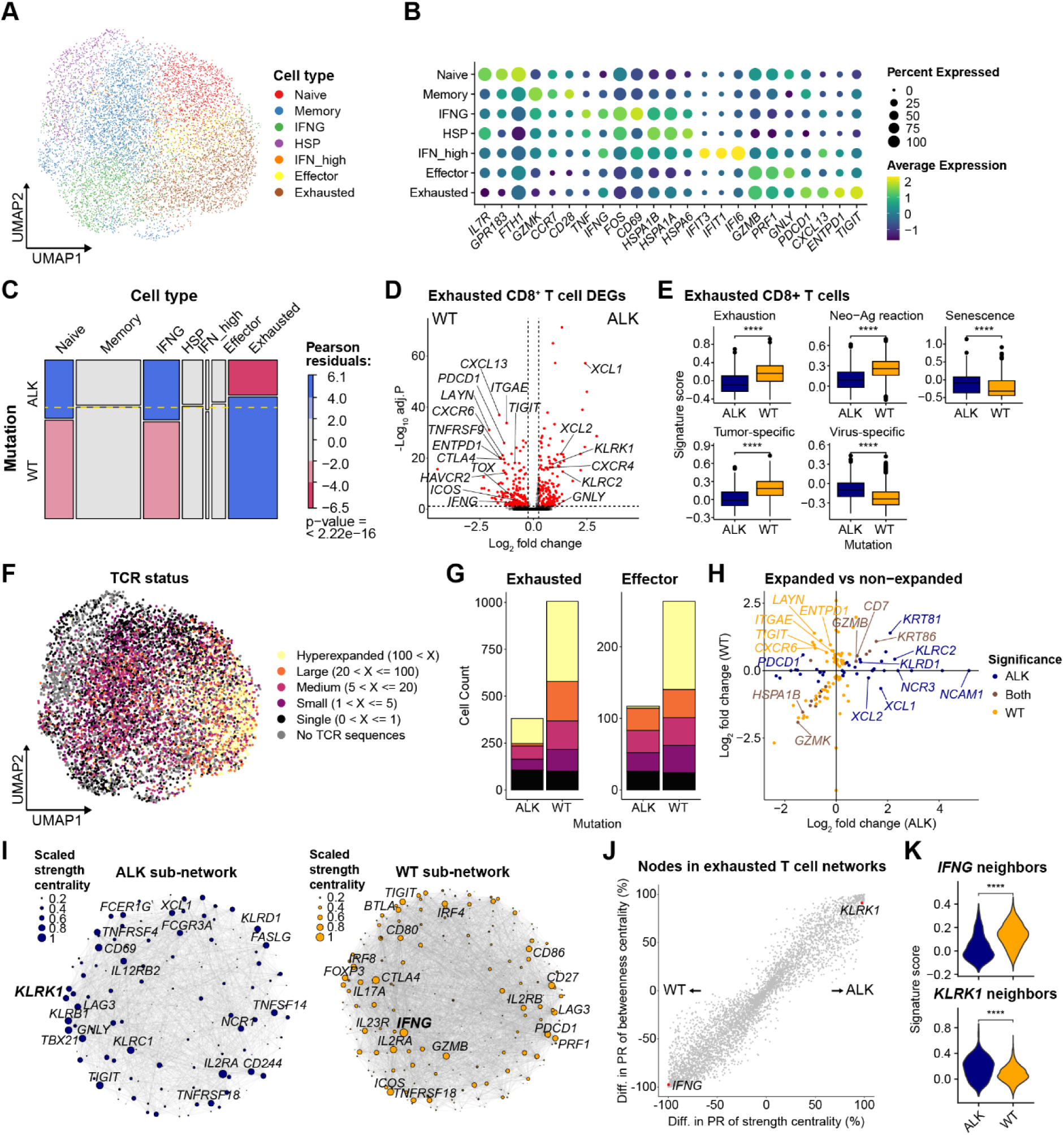
Dysfunctional states of CD8^+^ T cells in ALK-positive tumors. **A,** UMAP plot of CD8^+^ T cell subtypes. **B,** Dot plots of marker genes for each sub cell types of CD8^+^ T cells. **C,** Mosaic plots for compositional comparison of sub cell types of CD8^+^ T cells between WT and ALK-positive samples. The color indicates Pearson residuals, showing enrichment or depletion of the cell types for each mutation group. Significance was calculated using chi-square test. **D,** Volcano plot of DEGs of exhausted CD8^+^ T cells between ALK-positive and WT tumors. The p-values were calculated using the two-sided Wilcoxon rank-sum test with Bonferroni correction. The significant genes are colored red. **E,** Comparison of gene signature scores for gene sets related to functions and states of CD8^+^ T cells between two mutation groups. Significance of differences were calculated using the two-sided Wilcoxon rank-sum test. **F,** UMAP plot of CD8^+^ T cells colored for TCR-based clonal status. **G,** Bar plots for count of exhausted CD8^+^ T cells (left) and effector CD8^+^ T cells (right) for each clonal status in each mutation group. **H,** Visualization of two DEG analyses between expanded and non-expanded effector and exhausted CD8^+^ T cells for each mutation group. The genes are considered significant for each mutation group if the adjusted p-values < 0.05 (by two-sided Wilcoxon rank-sum tests with Bonferroni correction). **I,** ALK and WT sub-networks in sphere layouts from exhausted CD8^+^ T cells. Each node color and sizes are defined with the scaled strength centrality and top 20 genes are marked with gene names. **J,** Differences in percentile ranks of strength and betweenness centralities for each gene between exhausted CD8^+^ T cell networks from each mutation group. **K,** Gene signature scores for exhausted CD8^+^ T cells with neighboring nodes to *IFNG* (top) and *KLRK1* (bot) within WT and ALK group, respectively.

We compared the functional states of exhausted CD8^+^ T cells between ALK-positive and WT tumors using various T cell-related signatures^31–33^ including those for exhaustion, tumor neo-antigen reaction, senescence, tumor-specific, and virus-specific (bystander) T cells **(Supplementary Table 11)**. As expected, we observed enrichment of exhaustion, neo-antigen reactive, and tumor-specific functions in CD8^+^ T cells of WT tumors, while bystander and senescence-related functions were enriched in CD8^+^ T cells of ALK-positive tumors **(Fig. 5E)**. These results suggest that tumor-specific reactivity of CD8^+^ T cells is impaired in ALK-positive tumors potentially due to senescence. The resultant exhausted CD8^+^ T cells could explain the ineffectiveness of ICB treatment in ALK-positive patients.

For more in-depth analysis of antigen-specific T cell activation, we analyzed single-cell TCR sequencing datasets that matched our in-house single-cell RNA sequencing datasets. We observed highly expanded CD8^+^ T cells for effector and exhausted clusters **(Fig. 5F, Supplementary Fig. 5B)**, with fewer expanded cells and smaller TCR clonal sizes in ALK-positive tumors compared to WT tumors (**Fig. 5G**). These results indicate a lack of T cell antigenicity in ALK-positive tumors. We then performed DEG analysis between expanded and non-expanded effector and exhausted CD8^+^ T cells for each tumor group. The WT group showed an increase in exhaustion and tumor-reactivity-related genes, while the ALK-positive group exhibited decreased expression of these genes and increased expression of genes relevant to the NK receptors and NK marker genes **(Fig. 5H, Supplementary Table 12)**. These results reaffirm impaired tumor reactivity and more bystander-like function of CD8^+^ T cells in ALK-positive tumors.

Genes can alter their function through interactions with other genes. To investigate if changes in functional gene interactions underlie the reduced tumor-reactivity of CD8^+^ T cells in ALK-positive tumors, we constructed exhausted CD8^+^ T cell-specific networks for each mutation status using a method based on the reference interactome^34^ **(Supplementary Fig. 5C)**. We measured strength (weighted degree) and betweenness centrality for each network and compared percentile ranks between the two (**Fig. 5 I-J**). We identified T cell activation and exhaustion-related genes as hub genes, with *IFNG* as a top-ranked hub in the WT exhausted T cell network, aligning with its role in stimulating effector T cells^35^ and its known deficiency in ALK-positive patient^5^. Conversely, NK receptors like *KLRK1* had high centrality in the ALK-positive network, consistent with DEG analysis results. We re-evaluated hub genes based on neighboring gene expression in the network. *IFNG* neighboring genes had higher signature scores in exhausted T cells of WT tumors, while *KLRK1* neighboring genes were higher in exhausted T cells of ALK-positive tumors (**Fig. 5K**).

In summary, CD8^+^ T cells in ALK-positive patients showed lower tumor-reactivity and higher bystander-like signatures, explaining the ineffectiveness of immunotherapy and indicating noncanonical mechanisms for T cell dysfunction distinct from conventional T cell exhaustion.

### CD8^+^ T cells in ALK-positive tumors have altered gene regulatory network

T cell exhaustion is regulated at the epigenetic level by various genomic elements, including transcription factors (TFs) involved^36^. To gain novel insights into the dysfunctional states of CD8^+^ T cells caused by ALK rearrangement, we identified CD8^+^ T cell subtypes using scATAC-seq profiles and corresponding marker gene scores **(Fig. 6A-B, Supplementary Fig. 6A)**. These subtypes exhibited distinct chromatin accessibility patterns, with similar subtypes clustering together hierarchically **(Fig. 6C)**.

**Fig. 6.**
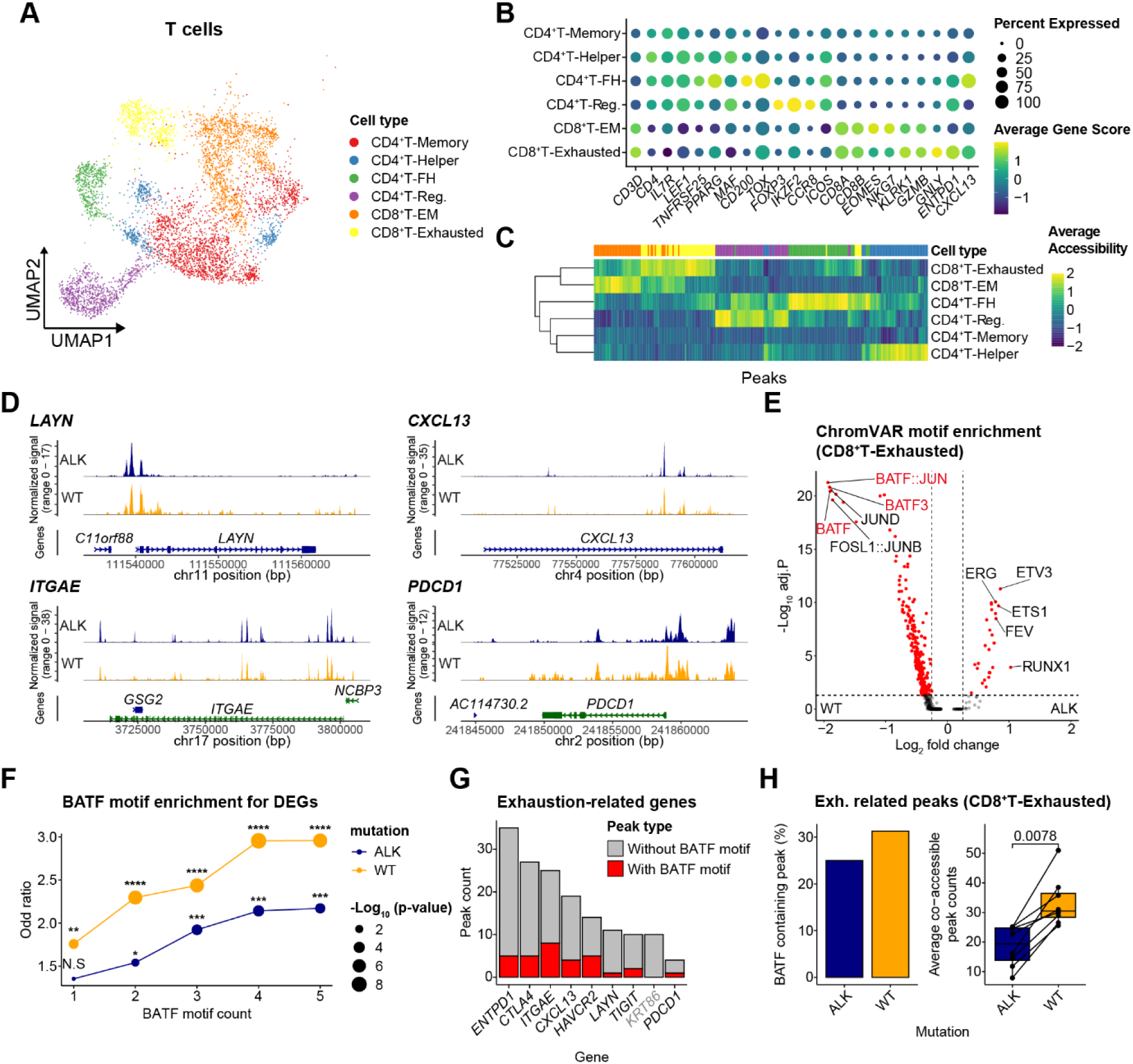
Changes in gene regulatory networks of CD8^+^ T cells in ALK-positive tumors. **A,** UMAP plot of T cells from scATAC-seq datasets, color-coded for sub cell types. **B,** Dot plot of marker genes for each sub cell types of T cells. **C,** Differentially accessible peaks for each sub cell types of T cells. Peak accessibilities were scaled column-wise and both columns and rows are clustered hierarchically. **D,** Genome tracks for *LAYN*, *CXCL13*, *ITGAE*, and *PDCD1*. Aggregated peak accessibilities for each mutation group were visualized along the chromosomes containing each gene. **E,** Volcano plot for differentially enriched chromVAR motifs for exhausted CD8^+^ T cells. The p-values were calculated using the two-sided Wilcoxon rank-sum tests with Bonferroni correction. **F,** Enrichment of genes with BATF motifs from DEGs of exhausted CD8^+^ T cells between ALK-positive and WT group. For each BATF motif count threshold, p-values and odd ratios were calculated using Fisher’s exact tests for each mutation group. **G,** Number of peaks related to exhaustion-related genes. Peaks were colored red for containing BATF motifs. **H,** Characterization of peaks from exhausted CD8^+^ T cells that are related to exhaustion genes. Percentages of peaks linked to the exhaustion genes that contained BATF motifs (Left). Average number of peaks co-accessible to peaks related to the exhaustion genes for each mutation group (Right).

Given the pronounced down-regulation of genes related to T cell exhaustion and tumor-reactivity in CD8^+^ T cells of ALK-positive tumors, we measured gene scores and chromatin accessibility for these genes using scATAC-seq analysis. Contrary to expectations, we did not observe clear depletion of accessibility for these genes in CD8^+^ T cells of ALK-positive tumors (**Fig. 6D, Supplementary Fig. 6B-C**). This suggests that while chromatin accessibility is similar, the TFs that are accessible might be different. We compared motif enrichments of TFs in exhausted CD8^+^ T cells between WT and ALK-positive groups (**Supplementary Table 13**). We found motifs of several BATF-related TFs enriched in the WT group, known for effector functions in CD8^+^ T cells^37^, while ETS1, required for NK cell differentiation^38^, was enriched in the ALK-positive group (**Fig. 6E**).

We hypothesized that BATF could be a key TF contributing to differences in T cell dysfunction. Using DEGs from scRNA-seq data analysis (**see Fig. 5D**), we measured whether WT group DEGs had more accessible peaks with BATF motifs. While ALK-positive group DEGs also had significantly more BATF motifs compared to non-DEGs, we observed stronger motif enrichment in WT group DEGs (**Fig. 6F**). All previously identified exhaustion and tumor-reactivity-related genes, except *KRT86*, had at least one peak with BATF motifs (**Fig. 6G**). Interestingly, *KRT86*, the only exhaustion-related gene upregulated in expanded cells from both WT and ALK-positive groups (see **Fig. 5H**), lacked BATF motif peaks. These findings suggest that while chromatin accessibility is similar, differences in TF activities, signified by their motif enrichment, may lead to differences in gene expression. Additionally, exhausted T cells of WT group had more nearby peaks containing BATF motifs with higher co-accessibility, indicating that these genes could be more co-regulated and connected to other genomic regions in WT group compared to the ALK-positive group (**Fig. 6H**).

## Discussion

Oncogenic driver mutations within the same cancer type introduce both new challenges and potential therapeutic avenues. While other major driver mutations such as EGFR have been extensively studied using various omics technologies at both bulk and single-cell resolutions, lung cancers with ALK-rearrangement are less studied, particularly at the single-cell level. Existing single-cell studies on ALK-positive lung cancer patients often suffer from low sample or cell counts. We aimed to leverage these less-studied samples, combined with our in-house datasets, for a comprehensively characterize ALK-positive lung cancers. This study provides valuable insights into the TME of ALK-positive lung cancers, potentially explaining recent failures in immunotherapy clinical trials for these patients.

Given that oncogenic mutations such as ALK rearrangement have a more direct impact on epithelial cells, we compared the characteristics of normal and malignant epithelial cells from ALK-positive and WT lung cancer tumor samples. ALK-positive epithelial cells exhibited higher stemness and more aggressive phenotypes, indicating greater cancer cell proliferation. Using external databases such as pan-cancer meta-programs^16^ or biological pathways, we identified evidence of aggressive cancer cells with stronger metastatic potential and immune interactions. Along the developmental trajectories of malignant cells, we observed an ALK-specific cell state resembling unfavorable tumor characteristics, supported by survival analysis using TCGA LUAD samples.

In broad immune landscape, we first observed an enrichment of innate immunity compartments and a depletion of adaptive immunity compartments, consistent with recent reports indicating immune-suppressive environments in ALK-positive tumors^5–7^. Among the immune cell types, myeloid cells showed largest increase in cell-cell interactions with malignant cells. Given that several subsets of myeloid cells, such as TAMs, can exert pro-tumoral immune-suppressive effects, we further dissected TAM sub-states. TAMs of ALK-positive tumors predominantly exhibited M2-like pro-tumoral phenotypes, with enriched sub-states linked to tumor-supportive activities such as lipid metabolism and adipogenesis. Additionally, TAMs of ALK-positive tumors tended to have stable states for pro-tumoral TAMs and transient states for anti-tumoral TAMs, supporting the existence of immune-suppressive TMEs.

Notable depletion of adaptive immunity in ALK-positive tumors potentially indicates a reduced capacity to sustain anti-tumoral immune function. The role of B cells in anti-tumoral activity has recently been highlighted in various types of cancers, including lung cancer. Reactivity of B cells to tumor antigens is critical for activating humoral response and TLS activity. We found that antigen reactivity and the capacity for TLS formation were reduced in B cells of ALK-positive tumors. Therefore, differences in compositional enrichment, activation status, and interactions among components of TLS-like structures indicated inadequate anti-tumoral responses by B cells of ALK-positive tumors, which could possibly lead to unfavorable TMEs for immunotherapy responses.

The most critical players in the adaptive immune response to tumors are likely CD8^+^ T cells. We found that exhausted CD8^+^ T cells were notably reduced in ALK-positive tumors, with downregulation of various exhaustion and tumor-reactivity-related genes. With low tumor-specificity and high bystander-like signatures, we hypothesized that CD8^+^ T cells of ALK-positive tumors exhibit distinct dysfunction mechanisms, such as senescence. Additionally, we observed decreased centrality for *IFNG* and increased centrality for NK receptor genes in the gene network specific to exhausted T cells, further indicating bystander-like phenotypes of exhausted CD8^+^ T cells in ALK-positive tumors. Chromatin accessibility profiles of CD8^+^ T cells yielded unexpected results, showing no noticeable differences in accessibility to the downregulated genes observed in gene expression profiles. Instead, we found fewer accessible peaks for BATF motifs in CD8^+^ T cells of ALK-positive tumors. Exhausted CD8^+^ T cells in WT tumors not only had more accessible BATF motifs but also higher co-accessibility to nearby peaks, indicating higher activity of trans-regulatory elements, leading to increased gene expressions.

In conclusion, we identified key cellular and molecular changes in ALK-positive tumor that could explain unfavorable immunotherapy response in ALK-positive lung cancer patients. In addition to the more aggressive traits of ALK-positive malignant cells, the immune-suppressive TAMs and depleted tumor-reactive activities in B cells and CD8^+^ T cells could account for the ineffectiveness of immune checkpoint blockades in patients with ALK-rearrangements. Further characterization of the distinct dysfunction mechanism in CD8^+^ T cells could enhance our understanding of adaptive immunity and potentially improve immunotherapy applications. Moreover, a careful examination of the epigenetic regulation of CD8^+^ T cells could open new avenues for reinvigorating T cells for improved tumor reactivity.

This study has several limitations. We integrated single-cell datasets from multiple cohorts, necessitating the integration of data from multiple sequencing platforms. While we believe our integration methods were carefully tested and successfully implemented, there may still be technical or cohort-specific biases. Additionally, including patients who have received immunotherapy would provide more direct evidence for our findings. Future studies with more single cell multi-omics datasets from patients before and after ICB treatments would further enhance our understanding of the oncogenic immune landscapes in lung cancer patients with ALK rearrangements.

## Methods

### Tumor specimen collection and dissociation

Patient samples were collected from individuals diagnosed with lung cancer who underwent surgical resection at Severance Hospital in Seoul, Republic of Korea, in 2020. A total of nine lung cancer patients were included in the study, of which three were ALK rearrangement-positive and 6 were wild-type (WT). Tumor tissues were finely minced and processed using a gentleMACS dissociator (Miltenyi Biotec, Gladbach Bergisch, Germany, Cat#130-093-235) followed by enzymatic digestion with the Human Tumor Dissociation Kit (Miltenyi Biotec, Cat#130-095-929) according to the manufacturer’s instructions. After a 1-hour incubation at 37°C, the samples were filtered through a 70-µm MACS SmartStrainer (Miltenyi Biotec, Cat# 130-098-462) into RPMI-1640 medium (Corning, Inc., Corning, NY, USA) supplemented with 10% fetal bovine serum (Biowest, Riverside, MO, USA) and then centrifuged at 300 ×g for 10 minutes. Tumor-infiltrating lymphocytes (TILs) were isolated using density gradient centrifugation with Ficoll (GE Healthcare). The collected cells were washed with RPMI-1640 medium and assessed for viability and cell counting using trypan blue.

### Oncogenic mutation analyses

EGFR mutation test was conducted using the PNAClamp EGFR Mutation Detection Kit (Panagene Inc., Daejeon, Korea). ALK rearrangements were identified using the VENTANA ALK (D5F3) CDx Assay (Ventana Medical Systems, Tucson, AZ, USA). For detecting ROS1 rearrangements, real-time PCR was performed with the ROS1 Gene Fusions Detection Kit (AmoyDx, Xiamen, China).

### Single-cell RNA and TCR sequencing

Libraries were generated using the Chromium controller in accordance with the 10x Chromium Next GEM Single Cell 5-v2_CellSurfaceProtein_UserGuide (CG000330). Cell suspensions were initially diluted with nuclease-free water to a target concentration of 10,000 cells. These suspensions were then combined with the master mix, Single Cell 5′ Gel Beads, and Partitioning Oil, and loaded into a Next GEM Chip K. Within the droplets, RNA transcripts from individual cells were barcoded and reverse-transcribed. The resulting cDNA was pooled and subjected to PCR enrichment. Following amplification, the cDNA was size-selected to generate three distinct library types: 5ʹ Gene Expression, V(D)J Enriched, and Cell Surface Protein. For the Cell Surface Protein Library, the cDNA pool underwent further enrichment using index PCR. The V(D)J Enriched Library required two rounds of amplification with specific primers. The 5’ Gene Expression Library was processed through end repair, addition of a single ‘A’ base, adapter ligation, and subsequent purification and PCR enrichment. Quantification of the purified libraries was performed using qPCR as outlined in the qPCR Quantification Protocol Guide (KAPA), and quality assessment was conducted using the Agilent Technologies 4200 TapeStation. Finally, sequencing was carried out on the HiSeq platform (Illumina) according to the read length specifications provided in the user guide.

### Single-cell ATAC sequencing and data preprocessing

Libraries were generated using the Chromium controller in accordance with the 10x Single Cell ATAC protocol (CG000168). Initially, nuclei were isolated and incubated with Transposase in a Transposition Mix. These transposed nuclei were then combined with master mix, Single Cell ATAC gel beads, and Partitioning Oil in a Chip E, where DNA fragments are barcoded in droplets during thermal incubation. The resulting barcoded DNA fragments were pooled and underwent sample index PCR. Quantification of the purified libraries was performed using qPCR as described in the qPCR Quantification Protocol Guide (KAPA), and quality assessment was conducted with the Agilent Technologies 4200 TapeStation. Finally, sequencing was carried out on the HiSeq platform (Illumina) following the read length specifications provided in the user guide.

### Single cell RNA seq processing and quality control

For the raw data from the Severance cohort, we utilized CellRanger^39^ (v6.1.1) with CellRanger GEX CRCh38 reference (2020-A) for read alignment, gene expression quantification, and cell calling. For the datasets from Wu et al.^8^ and Bischoff et al^10^., we used partially processed gene expression matrices for each sample. For the datasets from Maynard et al^9^., we aligned sequencing reads and quantified gene expression from the raw data with STAR^40^ (v2.7.9) and htseq^41^ (v2.0.4) with GRCh38 reference and Gencode v44. We used Seurat^42^ (v5.0.1) in R (v4.3.1) as a base during all processing and downstream tasks.

For removal of ambient RNA, we used CellBender^43^ (v0.3.0) with default parameters for the Severance cohort due to the availability of raw data from 10x Genomics platform and used decontX^44^ from Celda package (v1.16.1) for other datasets. After removing the ambient RNA, we filtered cells with mitochondrial gene percentage, read counts, and gene counts with adjusted thresholds for each cohort. We additionally removed doublets with DoubletFinder^45^ (v2.0.4) with 5% expected doublet rates.

### Single cell RNA seq integration and cell type annotation

As the initial step of multi-cohort integration, we matched gene aliases and available gene profiles. First, we converted all gene names into the consistent alias version by using ‘alias2Symbol’ function from limma^46^ (v3.56.2). If any genes became duplicated, we aggregated the expression from the duplicated genes into one. After gene symbol matching, we intersected lists of expressing genes from all four cohorts and removed any genes that did not exist for all four cohorts. When merging the datasets, we only considered those remaining intersected genes.

After merging, we initially used basic gene processing, dimension reductions, and clustering to identify cells from epithelial, stromal, and immune compartments. We used 3,000 variable features (filtered for ribosomal, TCR-related, and mitochondrial genes), and 50 PC dimensions for UMAP and clustering without major batch correction methods. After clustering, we annotated cells with canonical markers for epithelial, stromal, and immune cells along with reference-based cell type annotation results with Celltypist^47^ (v1.5.3) with its following databases; human lung atlas, immune all low, immune all high. We isolated each compartment and conducted major immune cell type annotations with immune cells.

With the identified immune cells, we repeated similar methods to identify cell clusters except for additional batch correction with scVI^48^ (v1.0.4). We selected 3,000 variable genes (filtered as before) and generated scVI model with sequencing platform and sample information as the covariates to account for multi-cohort integration. For subsequent UMAP visualization and clustering, we used 30 scVI latent representations. For the identification of major immune cell types, we used canonical immune markers for myeloid, CD4^+^ T, CD8^+^ T, NK, B, plasma, and mast cells along with the results from Celltypist. For sub-clustering analysis for each major immune cells, we used the same methods for re-clustering of identification of sub cell types.

### Single cell ATAC seq processing, quality control, and cell type annotation

After sequencing, we utilized CellRanger ATAC^49^ (v2.0.0) with CellRanger ARC GRCh38 reference (2020-A-2.0.0) for read alignment, identification of transposase cut sites and accessible peaks, and cell calling. Using the CellRanger ATAC outputs – “filtered_peak_bc_matrix.h5”, “singlecell.csv”, “peaks.bed”, and “fragments.tsv.gz” – each sample was initially processed with Seurat^42^ (v5.0.1) and Signac^50^ (v1.12.0) in R (v4.3.1). To reduce sample-specific effects, we generated a common set of peaks across all samples following a tutorial (https://stuartlab.org/signac/articles/merging). After generating the common peak set, we quantified quality-related metrics for each cell and filtered low quality cells. The cells with fewer than 2,000 reads, more than 5,000 reads, less than 15% of reads in the called peaks, more than four nucleosome signals (the ratio of mono-nucleosomal to nucleosome-free fragments), and less than 3 TSS enrichment (the ratio of fragments centered at the TSS to located in TSS-flanking regions). After the quality control, we merged all samples and conducted the initial dimension reduction with LSI. These dimensions were batch-corrected with harmony^51^ (v0.1.1) with sample information as variables. We visualized cells on UMAP coordinates calculated with 30 harmony-corrected LSI dimensions without the first dimension to reduce technical effects. We then identified clusters with ‘FindNeighbors’ and ‘FindClusters’ functions from Seurat with the same dimensions. For identification of cell types for each cluster, we used gene scores calculated with ‘GeneActivity’ function from Signac and used canonical marker genes. After isolating immune cells, we re-ran the same process except with 25 dimensions (also excluding the first dimension). After annotating immune cell clusters, we re-calculated cell type specific peaks with CallPeaks with immune clusters as input group. For further downstream analysis, we used these cell type specific peaks.

### Single-cell omics integration and label transfer from scRNA-seq to scATAC-seq profiled cells

To integrate cells from scRNA-seq and scATAC-seq, we used weighted nearest neighbor analysis from Seurat. With immune cells only, we first re-calculated gene scores for scATAC-seq datasets only using variable genes from scRNA-seq datasets. After normalizing and scaling gene scores, we generated transfer anchors with FindTransferAnchors with canonical correlation analysis reductions. We then generated RNA profiles for ATAC cells with ‘TransferData’ functions with gene expression matrix from scRNA-seq datasets and 25 harmony-fixed LSI dimensions (without the first dimension) for weight.reduction parameter. With the calculated RNA profiles for ATAC cells, we merged RNA cells and ATAC cells similar to merging two scRNA-seq objects. With the merged object, we used 25 PC dimensions for UMAP visualizations of both omics on the same dimensions. For cell type label transfer from scRNA-seq datasets to scATAC-seq datasets, we used ‘TransferData’ functions with immune cell type labels from scRNA-seq datasets and 25 harmony-fixed LSI dimensions (without the first dimension) for weight.reduction parameter.

### Cellular composition analysis

To investigate compositional differences in cell types between mutation groups, we used mosaic plots with VCD package^52^ (v1.4.12). The mosaic plots were colored with the Pearson residual which indicates deviations in the observed cell count for each group from the expected cell count. Blue color indicates that the observed cell counts were higher than expected and the red color indicates the opposite. The p-value for the goodness of fit for all subsets was calculated by the chi-square statistics. For the composition analysis of TAMs, we calculated percentages of TAMs in immune cells for each sample and grouped the proportions based on the mutation groups. For statistical significance, p-value was calculated with the two-sided Wilcoxon signed-rank sum test.

### Identification of malignant epithelial cells

For identification of malignant epithelial cells for both scRNA-seq and scATAC-seq datasets, we used the combinations of scATOMIC^53^ (v2.0.1) and scEVAN^54^ (v1.0.1). For scRNA-seq datasets, raw expression matrix from each sample was used as input for both tools with default parameters. For scATAC-seq datasets, we used raw gene score matrix instead. For high-confidential identification of malignancy, the cells were considered malignant when both tools identified them as tumor or malignant cells. While pan-cancer identification results from scATOMIC worked well for scRNA-seq, there were high numbers of seemingly mis-labeled cells for scATAC-seq datasets. Therefore, we used copy number variation-based identification of malignant cells through CopyKAT^55^ that was incorporated into scATOMIC.

### Assessing stemness of epithelial cells

To measure differentiation potential or stemness for normal and malignant epithelial cells, we used CytoTRACE^12^ (v0.3.3) which can predict developmental potential with the number of detectably expressed genes. We used ‘iCytoTRACE’ function with default parameters to measure CytoTRACE scores for multiple datasets. To compare the differences in CytoTRACE scores for each of normal and malignant epithelial cells between two mutation groups, we calculated *p*-values with the two-sided Wilcoxon signed-rank sum test.

### Differentially expressed gene (DEG) analysis

To identify genes or peaks with significant differences in expression between two groups, we used ‘FindMarkers’ function from Seurat. The genes or peaks were considered significant if they are expressed by at minimum 20% of cells with log_2_ fold change bigger than 0.25 and adjusted p-values lower than 0.01 unless otherwise stated. The *p*-values were calculated with the two-sided Wilcoxon signed-rank sum test and corrected for multiple tests through Bonferroni correction using all genes in the dataset.

### Pathway enrichment analysis

To discover enrichment of biological pathways for certain gene groups, we used EnrichR^56^ (v3.2) with their built-in databases. For each result, we annotated the databases that we selected. The *p*-values were calculated with the Fisher’s exact test and corrected using the Benjamini-Hochberg method. Gene ratio was calculated by dividing the number of intersected genes with input genes and genes in each pathway by total number of input genes.

### Gene signature scoring

To score each gene with gene sets or signatures of interests, we used ‘AddModuleScore’ function from Seurat package. In summary, the function calculates the average expression levels for each gene signature for each cell and subtracts the average expressions by the aggregated expression of control gene sets. The control gene sets are randomly selected from the genes within the same average expression bins.

### Cellular trajectory analysis

For generation of malignant cell trajectory, we used monocle2^57^ (v2.22.0). To identify genes for defining orders, we first subset genes that are expressed by more than 50 cells. We then grouped malignant cells into two groups based on whether their CytoTRACE scores were higher or lower than the median score. We then calculated differentially expressed genes between the two groups and selected top 1,000 genes according to their significance. During this process, we discovered that the malignant cells from P9 patient sample had exceptionally distinct gene profiles (with almost 0 percent of other cells expressing P9’s top DEGs) from other malignant cells with the third most cell counts, which were heavily affecting ordering genes and generating reserve flows on the trajectory. Therefore, we excluded malignant cells of P9 patient from the trajectory analysis. With the newly calculated top 1,000 ordered genes, we ran dimension reduction with DDRTree^58^ algorithm and ordered the cells with default parameters.

### Survival analysis with TCGA datasets

To identify whether the cells from each trajectory state would affect the clinical outcomes, we conducted survival analysis with bulk RNA sequencing data of LUAD patients from the TCGA cohort. We used clinical meta data for each patient and bulk expression matrix with transcripts per million quantifications. We identified DEGs of ALK-positive and WT malignant cells within the trajectory and DEGs of trajectory state 4 compared to state 1 and 5 using ‘FindMarkers’ function of Seurat. We selected genes passing q-value < 0.01 threshold. From these, we used the top 100 genes with the highest fold change and the top 100 genes with the lowest fold change. We used singscore^59^ (v1.14.0) to score enrichment of those DEG lists for each sample. With ‘simpleScore’ function of singscore, we used top 100 positive genes as upregulated genes (“upSet”) and top 100 negative genes as downregulated genes (“downSet”) to score each sample based on the difference between top 100 positive and negative DEGs. After dividing patients into two groups with high or low scores based on the median score, we measured the significance of differences in survival length with p-values from log-rank tests and hazard ratios from Cox regression. We visualized the result as Kaplan-Meier plot with survival^60^ (v3.2.13) and survminer (v0.4.9, https://github.com/kassambara/survminer) packages.

### Cell-Cell interaction (CCI) analysis

To measure interactions among different cell types and compare the interactions between mutation groups, we used CellChat^61^ (v2.1.1) with expression matrix for each mutation group following the default parameters. For CCI pairs, we subtracted interaction counts of WT samples from ALK-positive samples. For gene-gene pairs, we visualized interactions that are significant in at least one group with ‘netVisual_bubble’ function.

### RNA velocity and state transition analysis

To measure transitions among the macrophage states, we calculated RNA velocity with scVelo^62^ (v0.3.2). We only used single-cell RNA sequencing datasets from the Severance cohort due to requirement of having raw data. Using Velocyto^63^ (v0.17.17), we first quantified spliced and unspliced reads with CellRanger outputs. We then merged spliced and unspliced read matrix with the original RNA expression matrix for TAMs with the matching cell barcodes. We used scVelo with its dynamical mode in calculating RNA velocity. RNA velocity stream plots were visualized with ‘velocity_embedding_stream’ function from scVelo and the transition matrices with pair-wise cell transition probability values among all TAMs were extracted with ‘get_transition_matrix’ function. To calculate transition probabilities from one cell state to another cell state, we subset the transition matrix with rows from the first cell state (“From”) and with columns from the second cell type (“To”) and calculated the row-wise sums. To calculate significances of differences in distributions of such transition probability between two mutation groups, we used *p*-values calculated from Kolmogorov-Smirnov test.

### BCR analysis

For the Severance cohort datasets, we *in silico* reconstructed BCR sequences using TRUST4^64^ (v1.1.2). We utilized sorted BAM files from CellRanger, incorporating the default built-in hg38 BCR and IMGT references provided by TRUST4. Each B cell was annotated with its corresponding BCR sequences based on per-barcode reports, and clonal overlaps were assessed using scRepertoire^65^ (v2.0.0). Cells with BCR sequences that had two or more clones were classified as expanded.

### TCR analysis

With single-cell TCR sequencing raw data, we used CellRanger^39^ (v7.1.0) with CellRanger VDJ GRCh38 reference (alts-ensembl-7.1.0) for V(D) contig assembly, the contig annotation and CDR3 region location, and cell calling. Using the CellRanger VDJ outputs— filtered_contig_annotations.csv— and scReportoire^65^ (v2.0.0), we matched CD8^+^ T cells of scRNA-seq datasets from the Severance cohort with the same cells with VDJ profiles through the matched cell barcodes. We defined TCR status of each cell by counting the number of other cells with matching TCR clones and categorizing each cell into different count ranges: single (0 < count ≤ 1), small (1 < count ≤ 5), medium (5 < count ≤ 20), large (20 < count ≤ 100), and hyperexpanded (count > 100). For comparison between expanded and non-expanded cells, we defined the cells with TCR sequences having more than five clones as expanded.

### Cell-type-specific gene network analysis

To generate cell-type-specific gene networks for subtypes of CD8^+^ T cells of each mutation group, we used scHumanNet^34^ (v0.1.0). Among all cell-types-specific gene networks, we used the networks for exhausted CD8^+^ T cells of ALK-positive and WT tumors. For each network, we calculated betweenness and strength centrality using igraph^66^ (v1.5.1). For comparison of the network centralities, we measured percentile ranks for the centralities of genes and subtracted those of WT group from those of ALK-positive group. For visualization, we used DrL layout from igraph for the entire network and sphere layout for nodes within the same Louvain cluster as with *IFNG* for WT group and *KLRK1* for ALK-positive group. For sub-network plots, node sizes were determined by the strength centrality and only the top 20 nodes were annotated with their gene names.

### Motif enrichment analysis

To quantify the enrichment of TF motifs, we used chromVAR^67^ built in Signac^50^. First, using merged cell type specific peak sets from scATAC-seq, we added motif information from the JASPAR2020^68^ core vertebrate collection to the peaks. After adding motif information, we calculated chromVAR motif deviation score with ‘RunChromVAR’ function from Signac. With the chromVAR deviation matrix, we calculated differentially enriched chromVAR motifs with ‘FindMarkers’ function to compare exhausted CD8^+^ T cells between WT and ALK-positive samples. The fold-changes were calculated with the average difference in chromVAR deviation z-scores between two groups. We considered motifs significant if they exist in ≥10% of cells, with log_2_ fold change ≥ 0.25 and adjusted *p*-values < 0.05. The *p*-values were calculated with two-sided Wilcoxon rank-sum test and corrected for multiple tests with Bonferroni correction.

With the merged cell type specific peak sets that were annotated with motif information from scATAC-seq datasets, we calculated number of peaks with BATF motif for each gene in the DEG list. With different peak number threshold (from 1 to 5), we calculated significance of enrichment for BATF motif containing genes from DEGs of ALK-positive and WT samples compared to non-DEGs using the Fisher’s exact test. We visualized the results with p-values and odd ratios for each peak number threshold. Using the similar approach, we used the total number of peaks connected to exhaustion-related genes (*ENTPD1, CTLA4, ITGAE, CXCL13, HAVCR2, LAYN, TIGIT, KRT86, PDCD1*) and the number of peaks with BATF motifs to show that all genes except for *KRT86* contained peaks with BATF motif. To directly compare characteristics of peaks between exhausted CD8^+^ T cells of ALK-positive and WT samples, we calculated peaks linked to each gene using exhausted CD8^+^ T cells of ALK-positive or WT samples with ‘LinkPeaks’ function with default parameters. We then acquired linked peaks to the exhaustion-related genes for ALK-positive and WT samples. With these peak sets, we calculated proportions of peaks with BATF motif for each mutation group.

### Calculation of peak co-accessibility

To identify peaks that are co-accessible with their nearby peaks, we utilized Cicero^69^ built in Signac. For each mutation group, we subset exhausted CD8^+^ T cells using their scATAC-seq datasets. With ‘run_cicero’ function with default parameters, we constructed a list of co-accessible peaks for each mutation group. For each of the exhaustion-related genes, we calculated average number of peaks co-accessible to the peaks related to the gene for each mutation group. We then calculated paired Wilcoxon rank sum tests between two groups to measure the significance of differences in co-accessible peak counts.

## Data Availability

The single-cell RNA, TCR, and ATAC sequencing data generated in this study are deposited in the Gene Expression Omnibus database and will be available upon publication. The remaining data are available within the Article or Supplementary Information.

## Author contributions

H.R.K., and I.L. conceived the study. S.B. performed single-cell multi-omics data analysis under the supervision of I.L. E.S. assisted bioinformatic analysis. G.K. contributed sample preparation. H.R.K. and S.Y.P. organized clinical sample and data collections. C.Y.L. contributed to clinical sample collection. H.S.S. contributed to the pathological examination of tumor tissues. I.L. and H.R.K. contributed to the financial and administrative support for this study. S.B., H.R.K., and I.L. wrote the manuscript.

## Ethical approval

The studies were approved by the Institutional Review Board of Yonsei University Severance Hospital with IRB No 4-2018-1161-1. Written informed consent was obtained prior to enrollment and sample collection at Yonsei University Severance Hospital. The research conformed to the principles of the Helsinki Declaration.

## Acknowledgement

This research was supported by the Bio & Medical Technology Development Program of the National Research Foundation funded by the Ministry of Science and ICT (2021R1A2C2094629, 2017M3A9E9072669 to H.R.K, 2022M3A9F3016364, 2022R1A2C1092062 to I.L.). The work was supported in part by Brain Korea 21(BK21) FOUR program. This work was supported by the Technology Innovation Program (20022947) funded by the Ministry of Trade Industry & Energy (MOTIE, Korea). This work was supported by the Yonsei Fellow Program, funded by Lee Youn Jae.

## Competing interest statement

The authors declare that they have no conflicts of interest.

## Notes

### Competing Interest Statement

The authors have declared no competing interest.

## References

1 Siegel, R. L., Giaquinto, A. N. & Jemal, A. Cancer statistics, 2024. CA: a cancer journal for clinicians 74 (2024).

2 Chevallier, M., Borgeaud, M., Addeo, A. & Friedlaender, A. Oncogenic driver mutations in non-small cell lung cancer: Past, present and future. World Journal of Clinical Oncology 12, 217 (2021).

3 Du, X., Shao, Y., Qin, H. F., Tai, Y. H. & Gao, H. J. ALK-rearrangement in non-small-cell lung cancer (NSCLC). Thoracic cancer 9, 423–430 (2018).

4 Desai, A. & Lovly, C. M. Strategies to overcome resistance to ALK inhibitors in non-small cell lung cancer: a narrative review. Translational Lung Cancer Research 12, 615 (2023).

5 Schenk, E. L. Narrative review: immunotherapy in anaplastic lymphoma kinase (ALK)+ lung cancer—current status and future directions. Translational Lung Cancer Research 12, 322 (2023).

6 Wang, L. & Lui, V. W. Y. Emerging roles of ALK in immunity and insights for immunotherapy. Cancers 12, 426 (2020).

7 Guo, Y., Guo, H., Zhang, Y. & Cui, J. Anaplastic lymphoma kinase-special immunity and immunotherapy. Frontiers in Immunology 13, 908894 (2022).

8 Wu, F. et al. Single-cell profiling of tumor heterogeneity and the microenvironment in advanced non-small cell lung cancer. Nature communications 12, 2540 (2021).

9 Maynard, A. et al. Therapy-induced evolution of human lung cancer revealed by single-cell RNA sequencing. Cell 182, 1232–1251. e1222 (2020).

10 Bischoff, P. et al. Single-cell RNA sequencing reveals distinct tumor microenvironmental patterns in lung adenocarcinoma. Oncogene 40, 6748–6758 (2021).

11 Feitelson, M. A. et al. in Seminars in cancer biology. S25–S54 (Elsevier).

12 Gulati, G. S. et al. Single-cell transcriptional diversity is a hallmark of developmental potential. Science 367, 405–411 (2020).

13 Liberzon, A. et al. The Molecular Signatures Database (MSigDB) hallmark gene set collection. Cell Syst 1, 417–425 (2015). 10.1016/j.cels.2015.12.004

14 Fares, J., Fares, M. Y., Khachfe, H. H., Salhab, H. A. & Fares, Y. Molecular principles of metastasis: a hallmark of cancer revisited. Signal transduction and targeted therapy 5, 28 (2020).

15 Chen, Y., Ouyang, Y., Li, Z., Wang, X. & Ma, J. S100A8 and S100A9 in Cancer. Biochimica et Biophysica Acta (BBA)-Reviews on Cancer 1878, 188891 (2023).

16 Gavish, A. et al. Hallmarks of transcriptional intratumour heterogeneity across a thousand tumours. Nature 618, 598–606 (2023).

17 Network, C. G. A. R. Comprehensive molecular profiling of lung adenocarcinoma. Nature 511, 543 (2014).

18 Zhang, Q., Wang, X., Liu, Y., Xu, H. & Ye, C. Pan-cancer and single-cell analyses identify CD44 as an immunotherapy response predictor and regulating macrophage polarization and tumor progression in colorectal cancer. Frontiers in Oncology 14, 1380821 (2024).

19 Gillespie, M. et al. The reactome pathway knowledgebase 2022. Nucleic Acids Res 50, D687–D692 (2022). 10.1093/nar/gkab1028

20 Gene Ontology, C., et al. The Gene Ontology knowledgebase in 2023. Genetics 224 (2023). 10.1093/genetics/iyad031

21 Ma, R.-Y., Black, A. & Qian, B.-Z. Macrophage diversity in cancer revisited in the era of single-cell omics. Trends in immunology 43, 546–563 (2022).

22 Boutilier, A. J. & Elsawa, S. F. Macrophage polarization states in the tumor microenvironment. International journal of molecular sciences 22, 6995 (2021).

23 Chang, R. B. & Beatty, G. L. The interplay between innate and adaptive immunity in cancer shapes the productivity of cancer immunosurveillance. Journal of Leucocyte Biology 108, 363–376 (2020).

24 Waters, L. R., Ahsan, F. M., Wolf, D. M., Shirihai, O. & Teitell, M. A. Initial B cell activation induces metabolic reprogramming and mitochondrial remodeling. IScience 5, 99–109 (2018).

25 Yang, Y. et al. Pan-cancer single-cell dissection reveals phenotypically distinct B cell subtypes. Cell (2024).

26 Helmink, B. A. et al. B cells and tertiary lymphoid structures promote immunotherapy response. Nature 577, 549–555 (2020).

27 Raskov, H., Orhan, A., Christensen, J. P. & Gögenur, I. Cytotoxic CD8+ T cells in cancer and cancer immunotherapy. British journal of cancer 124, 359–367 (2021).

28 Hossain, M. A. et al. Reinvigorating exhausted CD8+ cytotoxic T lymphocytes in the tumor microenvironment and current strategies in cancer immunotherapy. Medicinal Research Reviews 41, 156–201 (2021).

29 Duhen, T. et al. Co-expression of CD39 and CD103 identifies tumor-reactive CD8 T cells in human solid tumors. Nature communications 9, 2724 (2018).

30 Kim, T.-S. & Shin, E.-C. The activation of bystander CD8+ T cells and their roles in viral infection. Experimental & molecular medicine 51, 1–9 (2019).

31 Wischnewski, V. et al. Phenotypic diversity of T cells in human primary and metastatic brain tumors revealed by multiomic interrogation. Nature Cancer 4, 908–924 (2023).

32 Chu, Y. et al. Pan-cancer T cell atlas links a cellular stress response state to immunotherapy resistance. Nature medicine 29, 1550–1562 (2023).

33 Oliveira, G. et al. Phenotype, specificity and avidity of antitumour CD8+ T cells in melanoma. Nature 596, 119–125 (2021).

34 Cha, J., Yu, J., Cho, J.-W., Hemberg, M. & Lee, I. scHumanNet: a single-cell network analysis platform for the study of cell-type specificity of disease genes. Nucleic acids research 51, e8–e8 (2023).

35 Jorgovanovic, D., Song, M., Wang, L. & Zhang, Y. Roles of IFN-γ in tumor progression and regression: a review. Biomarker research 8, 1–16 (2020).

36 Belk, J. A., Daniel, B. & Satpathy, A. T. Epigenetic regulation of T cell exhaustion. Nature immunology 23, 848–860 (2022).

37 Kurachi, M. et al. The transcription factor BATF operates as an essential differentiation checkpoint in early effector CD8+ T cells. Nature immunology 15, 373–383 (2014).

38 Taveirne, S. et al. The transcription factor ETS1 is an important regulator of human NK cell development and terminal differentiation. *Blood*, The Journal of the American Society of Hematology 136, 288–298 (2020).

39 Zheng, G. X. et al. Massively parallel digital transcriptional profiling of single cells. Nature communications 8, 14049 (2017).

40 Dobin, A. et al. STAR: ultrafast universal RNA-seq aligner. Bioinformatics 29, 15–21 (2013).

41 Putri, G. H., Anders, S., Pyl, P. T., Pimanda, J. E. & Zanini, F. Analysing high-throughput sequencing data in Python with HTSeq 2.0. Bioinformatics 38, 2943–2945 (2022).

42 Hao, Y. et al. Dictionary learning for integrative, multimodal and scalable single-cell analysis. Nature biotechnology 42, 293–304 (2024).

43 Fleming, S. J. et al. Unsupervised removal of systematic background noise from droplet-based single-cell experiments using CellBender. Nature methods 20, 1323–1335 (2023).

44 Yang, S. et al. Decontamination of ambient RNA in single-cell RNA-seq with DecontX. Genome biology 21, 1–15 (2020).

45 McGinnis, C. S., Murrow, L. M. & Gartner, Z. J. DoubletFinder: doublet detection in single-cell RNA sequencing data using artificial nearest neighbors. Cell systems 8, 329–337. e324 (2019).

46 Ritchie, M. E. et al. limma powers differential expression analyses for RNA-sequencing and microarray studies. Nucleic acids research 43, e47–e47 (2015).

47 Domínguez Conde, C., et al. Cross-tissue immune cell analysis reveals tissue-specific features in humans. Science 376, eabl5197 (2022).

48 Gayoso, A. et al. A Python library for probabilistic analysis of single-cell omics data. Nature biotechnology 40, 163–166 (2022).

49 Satpathy, A. T. et al. Massively parallel single-cell chromatin landscapes of human immune cell development and intratumoral T cell exhaustion. Nature biotechnology 37, 925–936 (2019).

50 Stuart, T., Srivastava, A., Madad, S., Lareau, C. A. & Satija, R. Single-cell chromatin state analysis with Signac. Nature methods 18, 1333–1341 (2021).

51 Korsunsky, I. et al. Fast, sensitive and accurate integration of single-cell data with Harmony. Nature methods 16, 1289–1296 (2019).

52 Friendly, M. & Meyer, D. Discrete data analysis with R: visualization and modeling techniques for categorical and count data. (CRC Press, 2015).

53 Nofech-Mozes, I., Soave, D., Awadalla, P. & Abelson, S. Pan-cancer classification of single cells in the tumour microenvironment. Nature Communications 14, 1615 (2023).

54 De Falco, A., Caruso, F., Su, X.-D., Iavarone, A. & Ceccarelli, M. A variational algorithm to detect the clonal copy number substructure of tumors from scRNA-seq data. Nature Communications 14, 1074 (2023).

55 Gao, R. et al. Delineating copy number and clonal substructure in human tumors from single-cell transcriptomes. Nature biotechnology 39, 599–608 (2021).

56 Kuleshov, M. V. et al. Enrichr: a comprehensive gene set enrichment analysis web server 2016 update. Nucleic acids research 44, W90–W97 (2016).

57 Qiu, X. et al. Single-cell mRNA quantification and differential analysis with Census. Nature methods 14, 309–315 (2017).

58 Mao, Q., Wang, L., Goodison, S. & Sun, Y. in Proceedings of the 21th ACM SIGKDD international conference on knowledge discovery and data mining. 765–774.

59 Foroutan, M. et al. Single sample scoring of molecular phenotypes. BMC bioinformatics 19, 1–10 (2018).

60 Therneau, T. M. & Lumley, T. Package ‘survival’. R Top Doc 128, 28–33 (2015).

61 Jin, S., Plikus, M. V. & Nie, Q. CellChat for systematic analysis of cell-cell communication from single-cell and spatially resolved transcriptomics. BioRxiv, 2023.2011.2005.565674 (2023).

62 Bergen, V., Lange, M., Peidli, S., Wolf, F. A. & Theis, F. J. Generalizing RNA velocity to transient cell states through dynamical modeling. Nature biotechnology 38, 1408–1414 (2020).

63 La Manno, G. et al. RNA velocity of single cells. Nature 560, 494–498 (2018).

64 Song, L. et al. TRUST4: immune repertoire reconstruction from bulk and single-cell RNA-seq data. Nature methods 18, 627–630 (2021).

65 Borcherding, N., Bormann, N. L. & Kraus, G. scRepertoire: An R-based toolkit for single-cell immune receptor analysis. F1000Research 9 (2020).

66 Csardi, M. G. Package ‘igraph’. Last accessed 3, 2013 (2013).

67 Schep, A. N., Wu, B., Buenrostro, J. D. & Greenleaf, W. J. chromVAR: inferring transcription-factor-associated accessibility from single-cell epigenomic data. Nature methods 14, 975–978 (2017).

68 Fornes, O. et al. JASPAR 2020: update of the open-access database of transcription factor binding profiles. Nucleic acids research 48, D87–D92 (2020).

69 Pliner, H. A. et al. Cicero predicts cis-regulatory DNA interactions from single-cell chromatin accessibility data. Molecular cell 71, 858–871. e858 (2018).

