## Supplementary Information for "Single-cell multi-omics reveals tumor microenvironment factors underlying poor immunotherapy responses in ALK-positive lung cancer"

### Contents


- ***Supplementary Figures & legends***
- ***Supplementary Table contents***

### Supplementary Figure

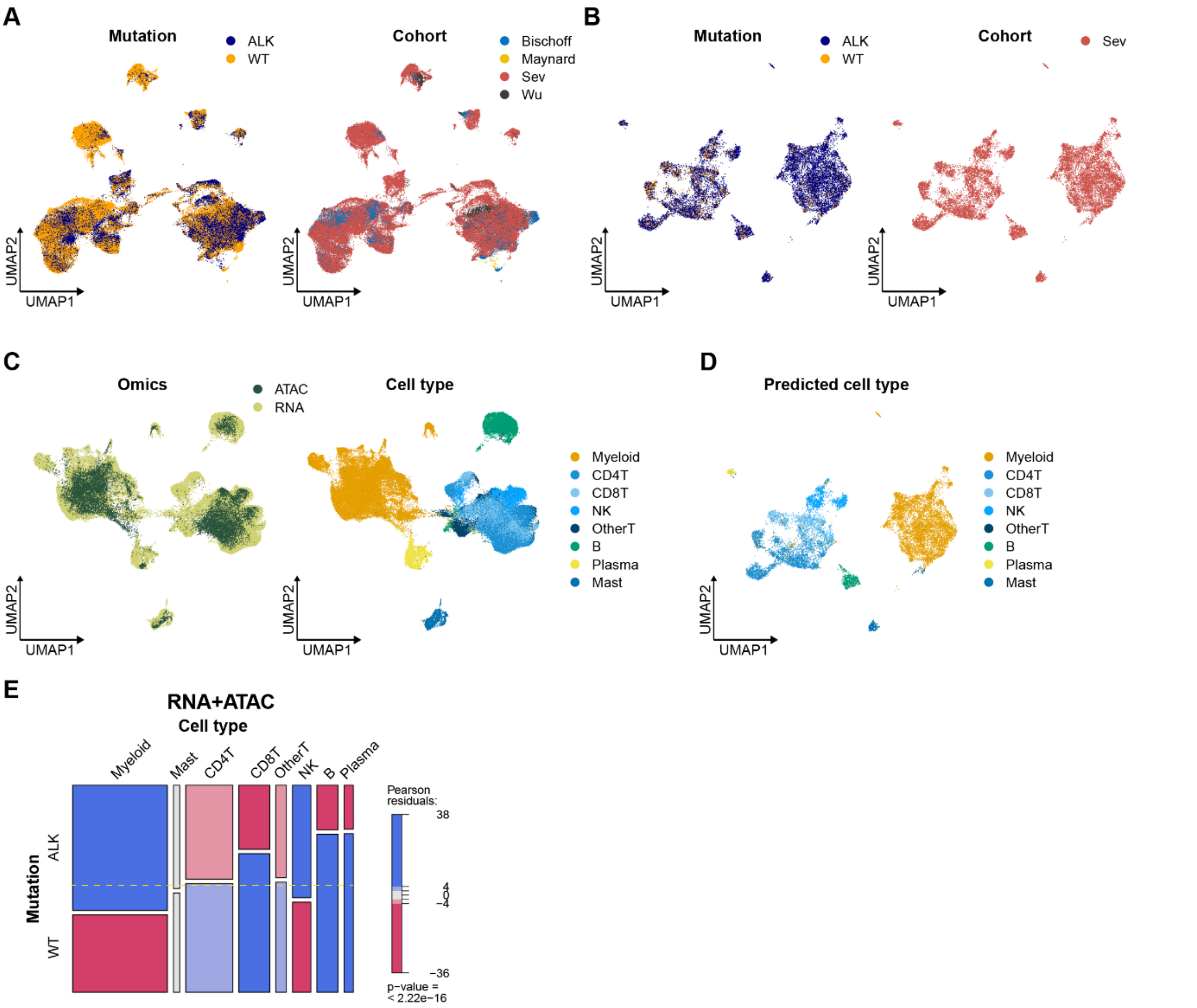


**Supplementary Fig. 1 | Integrative analysis of single-cell multi-omics data from this study. A,** UMAP plots of immune cells with integrated scRNA-seq profiles, color-coded for ALK rearrangement status (left) or cohorts (right). **B,** UMAP plots of immune cells with integrated scATAC-seq profiles, color-coded for ALK rearrangement status (left) or cohorts (right). **C,** UMAP plots of immune cells with integrated scRNA-seq and scATAC-seq profiles, color-coded for omics approach (left) or cell types (right). **D,** UMAP plot of immune cells with scATAC-seq profiles, color-coded for predicted cell types through label-transfer to scRNA-seq profiles. **E,** Mosaic plots for comparison of cell compositions between WT and ALK-positive groups with integrated multi-omics profiles. The color indicates Pearson residuals, showing enrichment or depletion of the cell types for each group. Significance was calculated using chi-square test.

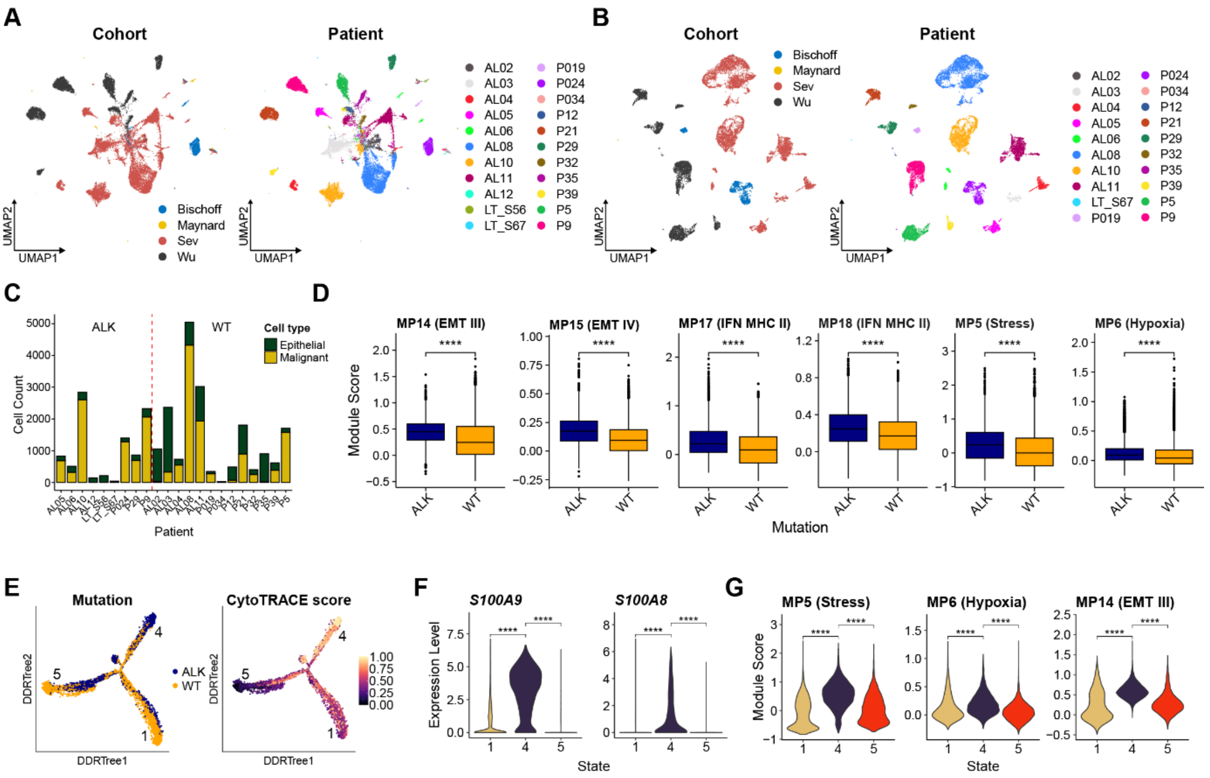


**Supplementary Fig. 2 | Overview of malignant cells in ALK-positive tumors. A,** UMAP plots of normal and malignant epithelial cells with single-cell RNA sequencing (scRNA-seq) profiles, color-coded for cohorts (left) and patients (right). **B,** UMAP plots of malignant epithelial cells with single-cell ATAC sequencing (scATAC-seq) profiles, color-coded for cohorts (left) and patients (right). **C,** Normal and malignant epithelial cells for each patient sample. **D,** Comparison of gene signature scores of the selected meta-programs for malignant cells between two mutation groups. Significance was calculated using the two-sided Wilcoxon rank-sum test. **E,** Trajectory plots of malignant cells with DDRTree algorithm, colored by mutation group (left) and CytoTRACE scores (right). **F,** Violin plots for gene expression of *S100A9* and *S100A8* for malignant cells in trajectory state 1, 4, and 5. Significance of differences was calculated using the two-sided Wilcoxon rank-sum test. **G,** Violin plots of gene signature scores for the selected cancer meta-programs for cells in trajectory state 1, 4, 5. Significance of differences was calculated using the two-sided Wilcoxon rank-sum test.

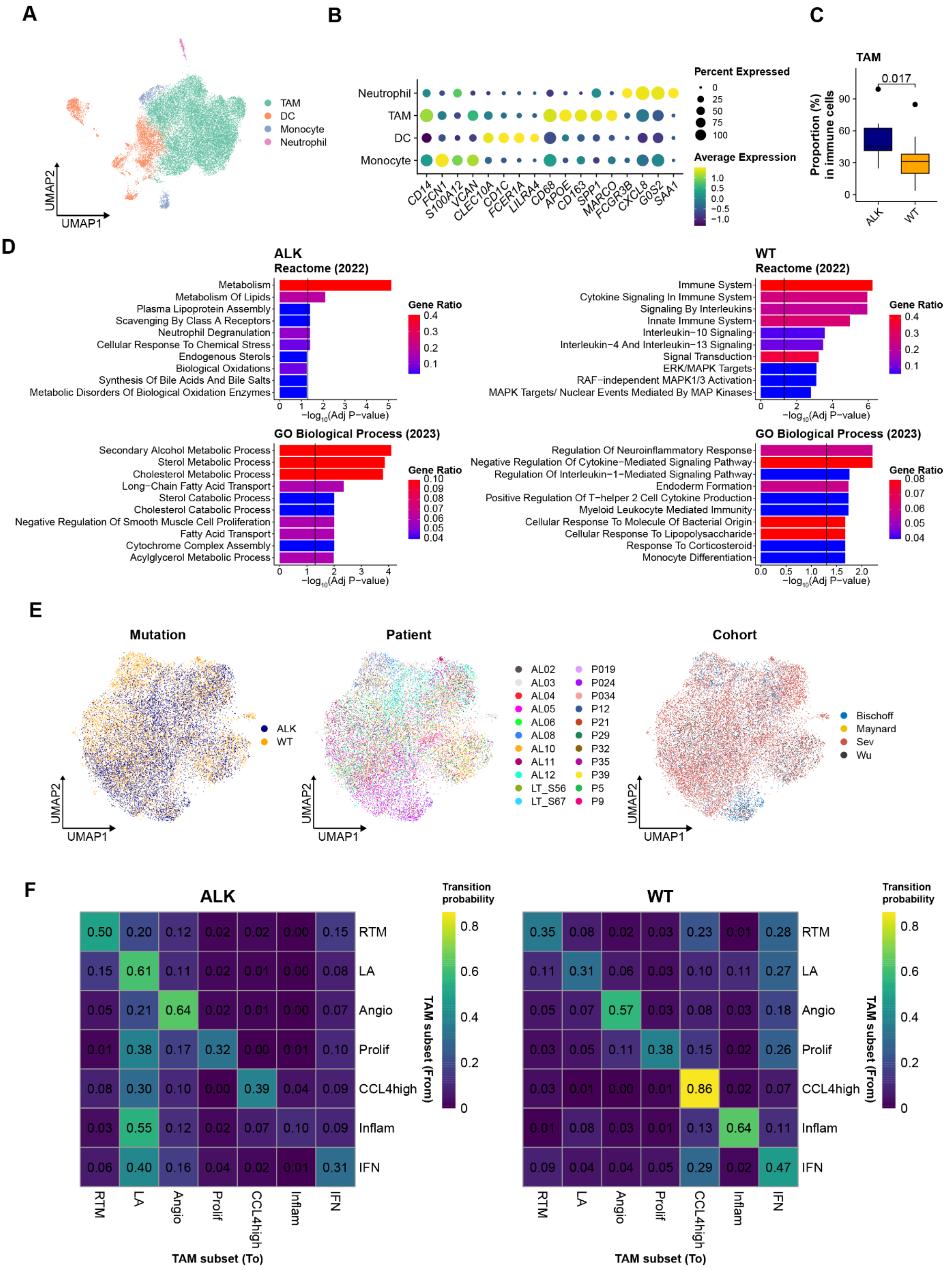


**Supplementary Fig. 3 | Myeloid cells and tumor-associated macrophages (TAMs) in ALK-**

**positive tumors. A,** UMAP plot of myeloid cells, color-coded for sub cell types. **B,** Dot plot of marker genes for each sub cell type of myeloid cells. **C,** Proportions of TAMs in immune cells for each mutation group. Significance of difference was calculated using the two-sided Wilcoxon rank-sum test. **D,** Enrichment of top 50 up-regulated genes in TAMs of ALK-positive tumors (left) and TAMs of WT tumors (right) for Reactome (2022) and GO biological process (2023) gene sets. The color indicates fractions of query genes that overlap with genes from the reference gene sets. The p-values was calculated using the Fisher's exact test with the Benjamini-Hochberg correction. **E,** UMAP plots of TAMs, color-coded for their mutation groups (left), patients (center), and cohorts (right). **F,** Heatmaps depicting cell type transitions from row to column among TAM subtypes from ALK-positive samples (left) and WT samples (right). The color indicates combined transition probabilities from one subset to another.

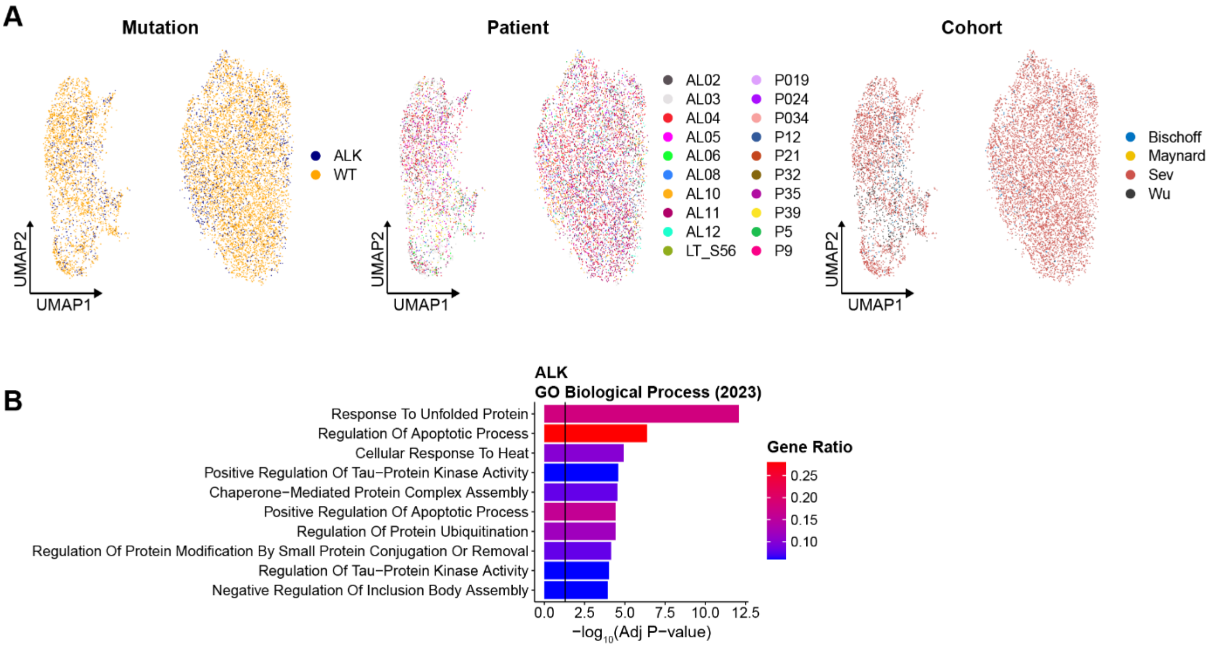


**Supplementary Fig. 4 | Overview of B cells in ALK-positive tumors. A,** UMAP plots of B cells, color-coded for mutation groups (left), patients (center), cohorts (right). **B**, Enrichment of top 50 up-regulated genes in memory B cells of ALK-positive tumors for GO biological process (2023) gene sets. The color indicates fractions of query genes that overlap with genes from the reference gene sets. The p-values was calculated using the Fisher's exact test with the Benjamini-Hochberg correction.

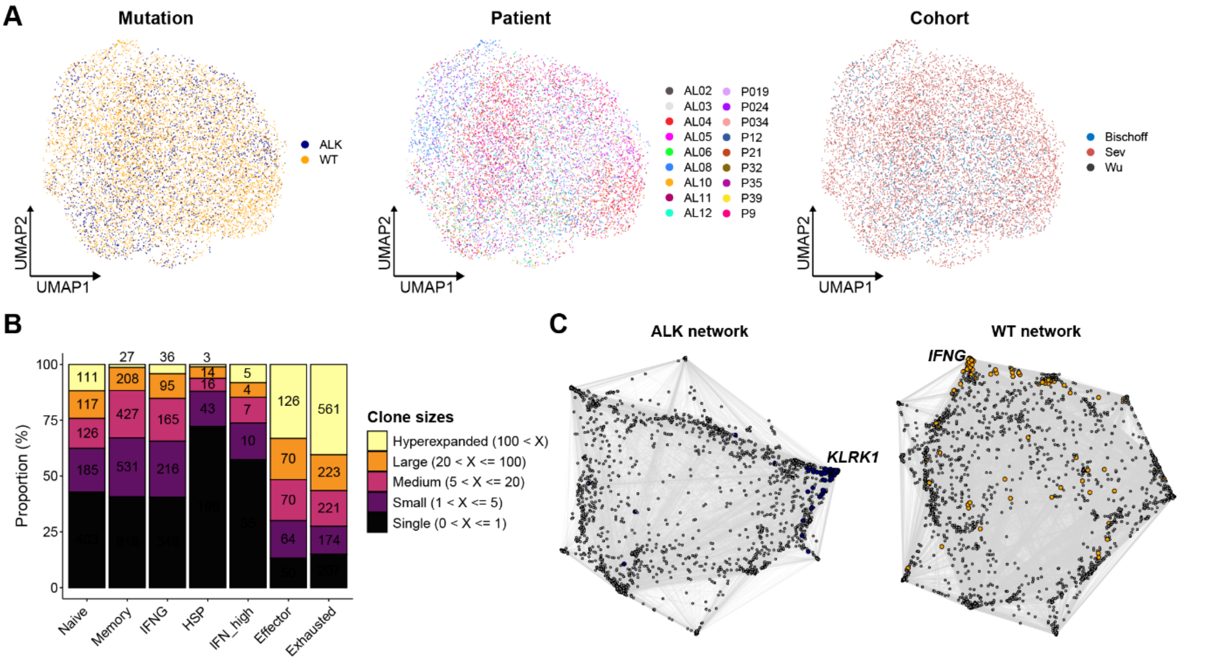


**Supplementary Fig. 5 | Single-cell RNA sequencing (scRNA-seq) data analysis for CD8$^{+}$ T cells. A,** UMAP plots of CD8$^{+}$ T cells, color-coded for their mutation group (left), patients (center), and cohorts (right). **B,** Proportions of CD8$^{+}$ T cells colored for their clonal status for each sub cell types. The numbers on the bars represent actual cell counts for each clonal status group. **C,** Gene network specific for CD8$^{+}$ T cells in ALK-positive and WT tumors with DrL network layout. The nodes included in the sub-networks of **Fig. 4I** for corresponding mutation group are colored navy for ALK-positive and orange for WT group.

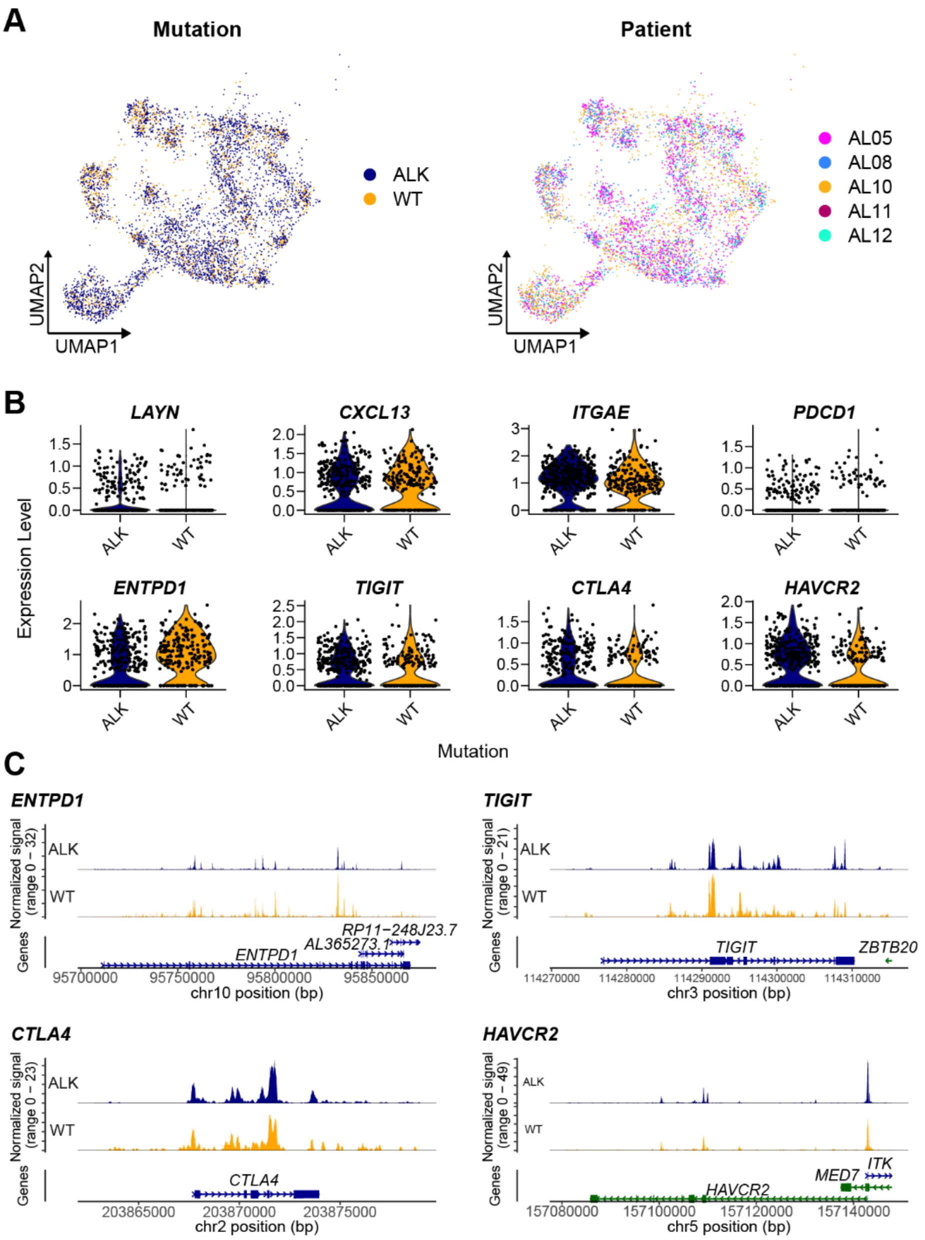


**Supplementary Fig. 6 | Single-cell ATAC sequencing (scATAC-seq) data analysis for CD8$^{+}$ T cells. A,** UMAP plots of T cells from scATAC-seq datasets colored for mutation groups (left) and patients (right). **B,** Violin plots of gene scores for exhausted CD8$^{+}$ T cells in each mutation group. **C,** Genome tracks for *ENTPD1*, *TIGIT*, *CTLA4*, and *HAVCR2*. Aggregated peak accessibilities for each mutation group were visualized along the chromosomes containing each gene.

### Supplementary Tables

**Supplementary Table 1 |** Sample information and single-cell data QC metrics

**Supplementary Table 2 |** Cell count information for samples and major cell types

**Supplementary Table 3 |** DEGs for each major immune cell type

**Supplementary Table 4 |** Top 100 up-regulated genes in ALK compared to WT malignant cells

**Supplementary Table 5 |** DEGs between ALK and WT samples and DEGs between state 4 and other states

**Supplementary Table 6 |** DEGs for myeloid cell subtypes

**Supplementary Table 7 |** Top 50 up-regulated genes for TAMs in ALK-postive tumors and those in WT tumors

**Supplementary Table 8 |** DEGs for TAM subsets

**Supplementary Table 9 |** DEGs for B cell subsets

**Supplementary Table 10 |** DEGs for $CD8^+$ T cell subsets

**Supplementary Table 11 |** Gene signatures for $CD8^+$ T cells

**Supplementary Table 12 |** Expanded vs non-expanded DEGs of effector and exhosted $CD8^+$ T cells

**Supplementary Table 13 |** ChromVAR DEGs
